# Within- and transgenerational plasticity evolve separately in a thermally adapted ectotherm

**DOI:** 10.64898/2026.11.18.767529

**Authors:** Corin Stansfield, Ana P.B. Costa, Matthew K. Brachmann, Gabriel Lawson-Duck, Bjarni Kristjánsson, Skúli Skúlason, Luca Lenzi, Sam Haldenby, Kevin Parsons

## Abstract

Climate change is imposing strong selection pressures on many wild populations, forcing them to adapt or face extinction. Conservation biology attempts to predict evolutionary responses to inform management but is hampered by a poor understanding of how variation arises. For many populations, responses to warming may rely substantially upon phenotypic plasticity, with cues from parental and offspring environments shaping development. However, how information from the environment, the genome, and the epigenome interacts to produce phenotypic variation is poorly understood. We investigated these factors, and their interactions, by using Icelandic populations of threespine stickleback (*Gasterosteus aculeatus*) that have adapted to geothermal warming. Using a split-temperature F_1_ to F_2_ hybrid cross, a novel morphometric approach, and reduced representation bisulphite sequencing, we elucidated the genetic architecture of transgenerational plasticity (TGP), within-generation plasticity (WGP), and that of morphological and methylation variation. This revealed that TGP and WGP have distinct genetic architectures involving different biological functions. The genetic architecture of traits was strongly influenced by both the parental and offspring thermal environments, potentially exposing different genetic variation to selection. Finally, we showed that TGP is favoured by selection and driven by the exposure of allelic variation by methylation. Our results suggested that predicting climate change responses should move beyond assessments of standing genetic variation to understand how TGP, WGP and the environment interact. More broadly, these findings improve our understanding of the processes that generate variation during adaptive divergence.

## Introduction

Climate change is a major driver of biodiversity loss, putting one third of species at risk of extinction (Urban, 2024). Effective conservation requires identifying both vulnerable populations and predicting their evolutionary responses. While some organisms can regulate physiology to buffer climatic shifts, ectotherms, representing ∼95% of species (Gilbert and Warner, 2026), are limited in this capacity, increasing vulnerability (Comte and Olden, 2017). In fish, elevated temperatures affect most phenotypes studied, including morphology, physiology, behaviour and life history (Alfonso et al., 2021; Campbell et al. 2021; Colchen et al., 2017; Crozier and Hutchings, 2014). Warming has been linked to reduced growth, fecundity, and survival (Devergne et al., 2023), suggesting direct negative fitness consequences. In particular, fish in polar regions are predicted to face warming rates 3-4 times greater than temperate regions (Pithan and Mauritsen, 2014; Previdi et al., 2021; Rantanen et al., 2022), while generally possessing narrow thermal tolerance ranges (Dahlke et al., 2020), increasing extinction risk. Understanding both the mechanisms that generate phenotypic variation and how selection acts on them is central to predicting evolutionary responses to climate change.

Phenotypic plasticity may drive adaptive phenotypic responses to climate change (Crozier and Hutchings, 2014). By allowing rapid responses to environmental cues within an individual’s lifetime ‘within-generation plasticity’ (WGP) is often implicated in persistence and adaptation to stressful environments (Schunter et al., 2018; Pilakouta et al., 2023; Potts et al., 2021) and has been shown to evolve in response to climate change related cues (Buckley and Kingsolver, 2025; Charmantier et al., 2008; Kingsolver and Buckley, 2017; Oyanoghafo et al., 2023). Environmental conditions experienced by a parent can also influence their offspring’s development in the form of transgenerational plasticity (TGP) (Harmon and Pfennig, 2021; Ryu et al., 2018). Inherited information, often passed on via epigenetic marks such as methylation (Herman and Sultan, 2016), can allow ‘tracking’ of environmental conditions, which is potentially adaptive if parental and offspring environments are similar (Galloway and Etterson, 2007; Wadgymar et al., 2018), or maladaptive if they differ (Herman et al., 2014; Spence-Jones et al., 2025). Indeed, parental exposure to elevated temperatures can reduce negative effects on offspring developing under the same conditions (Bernal et al., 2022; Donelson et al., 2012; Salinas and Munch, 2012; Shama et al., 2016; Spence-Jones et al., 2025), demonstrating a role for TGP in buffering environmental stress. Both TGP and WGP have been shown to have a robust genetic basis (Pfennig et al., 2010; Villicaña et al., 2021) and can therefore evolve (Nettle and Bateson, 2015; Pigliucci, 2005). Thus, by understanding how plasticity evolves, we can gain an understanding of how populations evolve mechanisms for short-term resilience under climate change.

How plasticity evolves depends on what heritable information is exposed to selection. In this sense, plastic traits may differ from non-plastic traits as environmental factors are influencing what heritable variation is expressed and exposed to selection (Lafuente and Beldade, 2019), shaping the evolution of climate change responses. TGP and WGP may then accelerate evolution by expressing plastic loci in relevant environments, allowing selection to connect environmental cues to adaptive phenotypes. In the context of TGP, genes that link parental environments to offspring phenotypes may be favoured and exposed to selection across multiple generations, facilitating evolution. Alternatively, loci may be ‘shielded’ from selection if environmental conditions are not met, allowing drift like dynamics to occur and limiting our ability to predict evolutionary responses (Gibson and Dworkin, 2004; Feiner et al., 2021; Paaby and Rockman, 2014). The evolution of plasticity will also depend on the relative contributions of TGP and WGP, including if and how they interact (Neylan et al., 2025). These two forms of plasticity may involve different molecular mechanisms (Bell and Hellmann, 2019; Jablonka and Raz, 2009; Kelly et al., 2012) and may be favoured or disfavoured under different evolutionary scenarios (Clement et al., 2023) or for different types of traits (Baduel et al., 2024; Déjeux et al., 2025). However, if TGP and WGP involve the same genes, and hence whether they can evolve separately to track environmental fluctuations is unknown. Thus, understanding the heritable variation underlying phenotypic plasticity, and how selection can act on these processes while avoiding constraints, is central to understanding the evolution of plasticity.

Insights into climate change responses can be gained from studying populations that have adapted to environmental stressors (O’Gorman et al., 2014). In this study, we leverage Icelandic populations of threespine stickleback (*Gasterosteus aculeatus*) that have diverged between adjacent geothermally heated and ambient habitats (Pilakouta et al., 2023). Previous work has shown heritable differences in morphology (Pilakouta et al., 2023; Smith et al., 2024) and metabolism (Brachmann et al., 2026) that aid survival in geothermal habitats.

Morphological divergence has occurred in just 75 years in some populations, suggesting adaptation occurring over timeframes relevant to current anthropogenic climate change (Pilakouta et al., 2023). Here, we investigated how TGP and WGP interact and evolve by constructing a hybrid cross with stickleback from geothermal and ambient environments, and rearing F_1_ and F_2_ generations at different temperatures in a full factorial experimental design. Using a novel morphometric approach, we identified morphological variation attributable to either TGP or WGP and performed QTL analysis to elucidate and compare the genetic architectures of both forms of plasticity. We predicted that i) TGP and WGP would involve distinct genes and biological functions, allowing the two forms of plasticity to evolve independently; ii) the genetic architecture of a trait, and hence what genetic information is exposed to selection, would be dependent on the current and past thermal environments; and iii) TGP will be favoured by selection for the ability to ‘forecast’ future environments and will be driven by methylation. By comparing both forms of plasticity within the same individual, and examining the underlying genetic and epigenetic architectures of trait variation, we gain unprecedented insights into the evolution of plastic responses to thermal conditions.

## Methods

### Animal husbandry and hybrid cross

Adult threespine stickleback (*Gasterosteus aculeatus)* were collected from adjacent geothermal (65.732260, -19.618937) and ambient (65.732191, -19.618574) habitats in northern Iceland, constituting the Sauðárkrókur (SKR) population pair (Pilakouta et al., 2023). The two habitats, separated by 30 metres, differ by up to 8°C during the summer, and have stickleback populations that have rapidly diverged between thermal habitats within the past 75 years (Pilakouta et al., 2023). These populations show significant genetic diversity and differentiation, despite their small size and geographic proximity (Brachmann et al., 2026). Sticklebacks were caught using unbaited minnow traps, kept in holding facilities at the University of Iceland at Hólar, and transported to aquarium facilities at the University of Glasgow, where they were kept as per Pilakouta et al. (2023). Fish were initially held under a 4:20 light-dark photoperiod for 6 weeks, followed by a gradual increase to a 24hr light regime that mirrored native conditions to induce spawning.

Our F_1_ hybrid intercross was established between a single geothermal male and ambient female using *in vitro* fertilisation (IVF) (Barber and Arnott 2000). Eggs were covered and left for 15 minutes to fertilise, before being split evenly between 12°C and 18°C. These water temperatures were chosen to simulate summertime geothermal and ambient habitats and have been found to induce phenotypic plasticity in morphology (Pilakouta et al., 2023). Fish (F_1_) from the same temperature treatment were crossed using IVF, with F_2_ progeny partitioned between 12°C and 18°C. This created four treatment groups with a tractable thermal history for two generations (Figure 1a); those that experienced 12°C for two generations (‘12_12’), 18°C for two generations (‘18_18’), fish whose parents experienced 12°C but themselves experienced 18°C (‘12_18’) and vice-versa (‘18_12’). A total of 294 F_2_ fish were reared to maturity across four treatment conditions before being euthanised with an overdose of benzocaine (10mg/L). We sampled fish from 18°C at ∼8 months of age and fish from 12°C at ∼10 months. This allowed us to compare fish of equivalent size, reducing the effects of allometry, while ensuring we were comparing individuals at equivalent developmental time points, as growth rates differ at the two temperatures (Pilakouta et al., 2023). Muscle tissue was taken from the right flank of each fish immediately post-euthanasia and stored in 100% ethanol at -80°C. Fish were then stored in 10% neutral buffered formalin.

**Figure 1.**
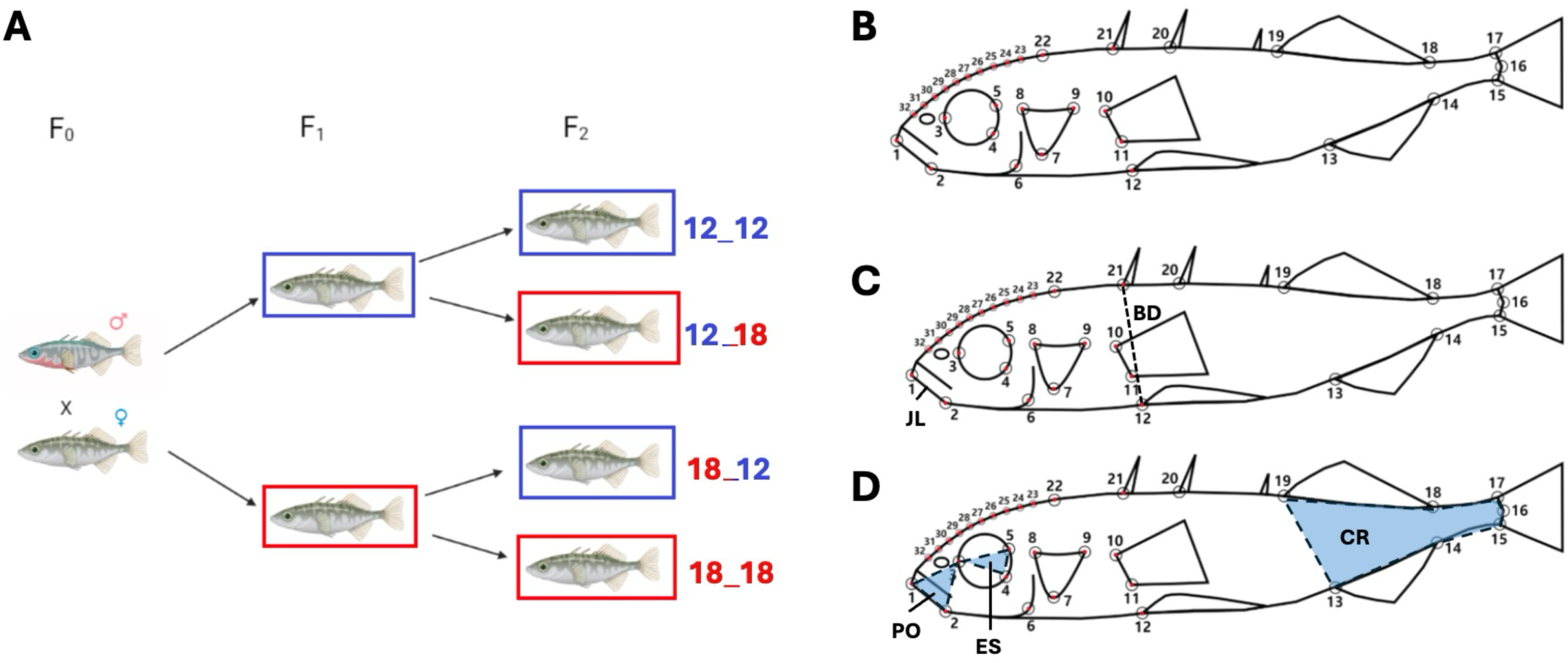
Hybrid intercross design and extraction of QTL traits. A) Hybrid intercross design. A geothermal (red) male and ambient (blue) female from the Sauðárkrókur (SKR) population pair were used to seed a hybrid cross. F1 progeny were split between 12°C (blue) and 18°C (red) and used to create an F2 generation which were similarly split across between 12°C and 18°C. Treatment groups were named in the format ‘12_12’ where the first number represents the F1 temperature while the second number represents the F2 temperature. **B) Landmarking scheme for geometric morphometrics.** Morphological traits were calculated from a set of 22 landmarks (large open circles) and 10 semi-landmarks (small open circles) placed in homologous locations. The anatomical landmarks were placed on the lower jaw (1, 2), eye orbit (3‒5), preopercle (6), opercle (7‒9), pectoral fin insertion points (10, 11), anterior tip of pelvic spine (12), anal fin insertion points (13, 14), caudal fin insertion points (15, 17), caudal border of the hypural plate at the lateral midline (16), dorsal fin insertion points (18, 19), dorsal spine insertion points (20, 21), and the posterior tip of the frontal bone (22). 10 semi-landmarks (23-32) were placed evenly between the posterior tip of the frontal bone (23) and the anterior tip of the upper jaw (32). ‘Head’ landmarks are indicated with a red dot. ‘Body’ landmarks are those lacking this red dot, apart from landmarks 12 and 21 which are included in both subsets. **C) Linear traits.** Linear measurements of anatomical features. JL = Jaw length; BD = Body Depth. **D) Area-based traits.** Areas of anatomical features are shaded in blue. PO = Pre-Orbital; ES = Eye Size; CR = Caudal Region.

### Genotyping and linkage map construction

DNA was extracted using Qiagen DNeasy blood and tissue kit following the manufacturer’s instructions. Sex was determined via the amplification of sex specific Y-chromosome insertions, visualised on an agarose gel as per Bakker et al. (2017). Genotype-by-sequencing (GBS) library preparation and SNP calling was conducted as per Brachmann et al. (2026), with the exception that the sex chromosome (19) was retained. Briefly, DNA was digested with ApeKI and ligated to custom adapters to construct genotype-by-sequencing libraries, which were sequenced with a read length of 150bp on an Illumina HiSeq 4000 sequencer at the Center for Genomic Research, University of Liverpool. Reads were aligned to the V5 stickleback reference genome (Nath et al., 2021) and SNP calling was performed using ANGSD v0.929 (Korneliussen et al., 2014). Sites with a minor allele frequency (MAF < 0.05) or were present in less than 80% of individuals were dropped. See Brachmann et al. (2026) for full details.

Markers that did not map to any chromosome were dropped, as were those mapping to the male-specific Y-chromosome. The remaining 398k markers were used to construct a linkage map based on recombination frequencies using Lep-Map3 (Rastas, 2017). Parental genotypes were called and uninformative markers (e.g. homozygous in both parents) were removed using the *ParentCall2* module. Data were cleaned using the *Filtering2* module to remove markers segregating in a non-mendelian fashion or missing in more than 10% of individuals, leaving 78k markers. Linkage group assignment was performed using the *SeparateChromosomes2* module, which assigned markers to linkage groups by computing all pairwise logarithm of odds (LOD) scores between markers and joined pairs with a LOD score higher than a user-defined value, in this case 16. These parameters were chosen as they produced at least one linkage group containing markers that mapped exclusively to one of the 21 stickleback chromosomes. Distances between markers were calculated with the *OrderMarkers2* module using a given order determined from the physical position of each marker aligned against the V5 genome (Nath et al., 2021).

### Geometric morphometrics and the isolation of WGP and TGP variation

To quantify morphological variation, F_2_ fish were bleached to remove skin pigmentation (Potthoff, 1984) and stained with alizarin red S to highlight bone morphology, before being washed with 75% glycerol to remove excess stain. Specimens were positioned and photographed on their left-hand side using a Canon EOS 1100D digital camera (Canon inc., Tokyo, Japan) with an EF-S 18-55mm lense set at a fixed height and angle using a copy stand (Pilakouta et al., 2023), and a ruler in each photo to provide scale. Landmarks were placed at 22 homologous locations using TPSdig (Rohlff, 2017) as per Pilakouta et al. (2023) (Figure 1B). 10 semi-landmarks were placed equally spaced from the anterior tip of the upper jaw to the posterior tip of the frontal bone to capture curvature of the head. Data were loaded into R version 4.3.1 (R Core team, 2023) where subsequent morphometrics were performed primarily using the ‘geomorph*’* package v4.0.7 (Adams and Otárola-Castillo, 2013).

The effects of scale, rotation, and translation were removed from our landmark data through a Generalized Procrustes Analysis using the *gpagen* function. We then leveraged our experimental design to isolate morphological variation attributable to either the parental temperature (TGP) or offspring temperature (WGP). Briefly, Procrustes-corrected landmarks were regressed onto a factor representing either the F_1_ or F_2_ temperature using the *ProcD.lm* function. This generated a ‘fitted value’ in the form of a set of landmark co-ordinates for each temperature. For a fish reared at 12°C, we would take their landmark co-ordinates and ‘subtract’ the fitted value for 18°C (and vice-versa). Thus, the difference between a fish’s morphology and the fitted value for the alternative temperature was attributed to phenotypic plasticity from that generation. This was repeated for all F_2_ fish, before adding the consensus shape for all fish to each set of landmarks, creating landmark data comparable in structure to the raw data and for which all morphological differences could be attributed to the focal generation. Potential interactions between F_1_ and F_2_ temperature were not included in this process but were controlled for at the quantitative trait loci (QTL) modelling stage. This approach left us with three sets of landmarks; one representing unmodified morphological variation (Procrustes-corrected landmarks), one representing variation attributable to TGP and another attributable to WGP. Our approach was akin to removing the effects of allometry from morphometric data in that fitted values (variation attributed to allometry) were excluded, and residuals (allometry-free variation) were taken forward for analyses. Here, we simply reversed this logic and took the variation attributed to the explanatory variable (F_1_ or F_2_ temperature) forward for analyses. All subsequent morphometrics and analyses were performed on each set of landmarks individually.

Variation in morphology, TGP, and WGP was quantified in a set of morphological traits highlighted previously in this system (Pilakouta et al., 2023; Smith et al., 2024) and other stickleback populations (Peichel and Marques, 2017). Traits were chosen that underlie heritable variation in head and body shape adaptively diverging between geothermal and ambient ecotypes (Pilakouta et al., 2023; Smith et al., 2024), and which may show TGP and WGP (Supplementary Table 1). These firstly consisted of linear measures (jaw length and body depth; Figure 1C) calculated using the *interlmkdist* function. These traits have been shown to effect foraging (McGee et al., 2013) and swimming efficiency (Langerhans, 2008) respectively and are divergent between geothermal and ambient ecotypes (Pilakouta et al., 2023). Next were several measures of the area of anatomical features (Figure 1D) calculated by partitioning a subset of landmarks into a geomorph data frame and extracting centroid size. The pre-orbital region has been identified as an evolvable module facilitating trophic diversification in perciformes (Cooper et al., 2010; Parsons et al., 2011). Eye size relates to the survival of stickleback in geothermal and ambient habitats (Smith et al., 2024) while the dimensions of the caudal region effects manoeuvrability in different microhabitats (Langerhans, 2008). Linear measurements and areas were regressed against log centroid size, with standardised residuals taken forward as traits for QTL analysis. Trait values were calculated using homologous landmarks from each of the three landmark sets to capture variation in unaltered morphology, TGP, and WGP sets respectively.

Finally, we partitioned landmarks and semi-landmarks into ‘head’ and ‘body’ landmark sets (Figure 1B) and quantified morphological variation in each that was attributable to TGP and WGP. Firstly, relative warps scores were calculated from landmark sets using the program tpsRelW32 v1.49 (Rohlf, 2010). Relative warp scores represent orthogonal axes of multivariate shape variation and are equivalent to principal components (Zelditch et al. 2012). Variation attributable to F_1_ and F_2_ temperature respectively was then isolated using discriminant function analyses, run using the *candisc* function in the “candisc” package (Friendly and Fox, 2017). This package allowed us to isolate variation in scores attributable to each explanatory variable (generation), while controlling for effects of the other. Hence, two models were run, extracting variation in shape attributable to F_1_ and F_2_ temperature respectively, with the first canonical root score for each individual being used for QTL analysis. This was performed separately for ‘head’ and ‘body’ landmark sets.

### QTL analysis

The genetic architecture of unaltered morphological variation, as well as that attributable to TGP and WGP were investigated using a QTL approach. QTL analysis was run on each trait and data type (Morphology, TGP, WGP) sequentially. We first performed a statistically liberal scan to find suggestive QTL with a LOD score greater than a 95% threshold determined by 1,000 permutations for each model, using the *scanone* function in the “qtl” package (Broman and Sen, 2009). Chromosomes containing a marker exceeding significance, or within 1 LOD of the significance threshold, were taken forward as putative candidates for subsequent refinement. We then performed multiple QTL mapping (MQM) for these putative candidates using the *fitqtl* function, using combinations of suggestive loci as cofactors to further validate our models. Sex was coded as a categorical variable for all models. When modelling TGP variation for a trait, the F_2_ temperature was included as a cofactor, and vice-versa for WGP, to control for any interaction between F_1_ and F_2_ temperatures. Multiple peaks on a single linkage group were treated as distinct QTL if the LOD value dropped by 2 between the putative peaks, akin to the 2-LOD drop widely used in QTL mapping (MacDonald et al., 2022; Van Ooijen, 1999). 1.5-LOD support intervals were calculated for each peak using the *lodint* function. The percentage of variation explained (PVE) was calculated using the formula:

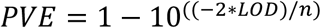

Where *n* is the number of individuals in the hybrid cross. To avoid biases caused by replicate QTL in potentially colinear traits (as highlighted by Miller et al. (2014)) we adopted the approach of Erickson et al. (2016) by splitting our traits into three classes -head morphology, body morphology and methylation (Supplementary table 1). QTL were excluded whose confidence intervals overlapped with another QTL within that trait class and model type, with the QTL with the higher PVE kept in the case of conflicts. This resulted in a set of 51 QTL (10 Morphology, 26 TGP, 15 WGP) that were taken forward for subsequent analyses.

### Genetic similarity of TGP and WGP

TGP and WGP were predicted to be genetically distinct, potentially allowing them to evolve independently. This was investigated by testing the similarity of their genetic architecture at various biological scales. Similarity of the genetic architectures of TGP and WGP was measured by comparing how explanatory power was distributed across the genome. LOD score profiles were extracted from *scanone* models for TGP and WGP of each morphological trait and partial pairwise correlation scores were calculated using the *pcor* function in the “ppcor” package (Kim, 2015). Whether TGP and WGP mapped to different regions of the genome was tested by counting instances where 1.5-LOD confidence intervals for QTL implicated in TGP and WGP overlapped. Finally, the tendency of TGP and WGP QTL to cluster on different chromosomes was tested by running partial pairwise correlation analysis as above comparing the distribution of TGP and WGP QTL across all chromosomes.

### Identification of underlying genes and biological functions

Genes and biological processes underpinning TGP and WGP were investigated through fine-scale QTL mapping and gene ontology (GO) analysis. QTL models containing multiple peaks can return large confidence intervals spanning much of the chromosome, even if this includes regions with low LOD scores (Broman and Sen, 2009). We therefore narrowed our search for candidate genes in regions i) falling within 1.5-LOD confidence intervals and ii) where LOD ≥ 3. By combining two measures of significance, we focused our search to regions of the genome that were more likely to be causative, identifying what we refer to as ‘candidate regions’. Chromosome walks were performed by looping ANOVAs for every marker in the raw GBS data, including those not included on the final linkage map, falling within the candidate region against the focal trait, allowing genotype-phenotype correlations to be mapped at greater resolution. All markers with a R^2^ value above 0.05 were taken forward, as this approximately corresponded to the top 5% of scores across models. Biological functions of any gene containing such a marker were investigated by performing a database search on NIH (https://www.ncbi.nlm.nih.gov/gene/), with genes with biological functions potentially relating to the trait of interest being recorded as putative candidates. Whole genome data from 14 geothermal and 14 ambient stickleback were used to measure selection on candidate genes in the wild. F_st_ scores were calculated per locus between geothermal and ambient ecotypes using plink (version 1.9) (Purcell et al., 2007; Chang et al., 2015) as outlined in Brachmann et al., (2026). A final list of candidate genes was created by filtering only those genes containing a locus with an F_st_ score above 0.05.

Biological functions underlying morphological variation and plasticity were identified using gene ontology (GO) analysis. The ensembl IDs for all genes (n=3188) that mapped to a candidate region on the V5 genome (Nath et al., 2021) were extracted. GO analysis was performed using metascape (Zhou et al., 2019), using gene IDs for the most closely related species to threespine stickleback, *Danio rerio,* for data input and analysis. Firstly, gene lists pertaining to the different model types were submitted separately (TGP=2122; WGP=1066). We then compared similarity in terms enriched for TGP and WGP by submitting lists of gene IDs from both model types and running GO analysis simultaneously using the multiple gene lists option.

### Quantifying the effect of temperature on genetic architecture

Whether the cumulative effect of within-and transgenerational thermal environments influenced the genetic architecture of a trait was tested by dividing our data into the four multi-generational treatment groups (12_12, 18_18, 12_18, 18_12) and running unaltered morphology QTL models for each treatment group in turn. A LOD threshold of three was used to identify QTL and candidate regions across models and treatment groups. Similarity of LOD distributions and GO terms was compared as above. Finally, we tested whether inheritance patterns were affected by parental and offspring temperatures. Allele effects were quantified by calculating the difference in mean phenotypes between individuals with each homozygous genotype, known as the homozygous substitution effect (HSE). This was repeated for all four treatment groups and all resulting morphology QTL.

### Methylation data collection and DMC determination

To assess patterns of methylation data we used reduced representation bisulfite sequencing (RRBS) using flank tissue from our F_2_ fish. While providing an environmentally induced molecular phenotype this also allowed us to quantify underlying genetic variation and characterise interactions with TGP and WGP. To begin, 150ng of gDNA was digested with the MspI restriction enzyme, followed by end repair using an NEBNext ® Ultra ™ II DNA Library Prep Kit for Illumina ® (New England BioLabs ® Inc.). Adapter ligation was performed with the NEBNext ® Ultra ™ II kit and NEBNext ® Multiplex Oligos for Illumina ® (Methylated Adaptor, Index Primers Set 1). The DNA purification and size selection (keeping fragments of 100-400 bp) were performed with AMPure XP beads. Purified, adapter-ligated DNA was bisulfite treated using EpiMark ® Bisulfite Conversion Kit (New England BioLabs ® Inc.). PCR was performed on bisulfite converted DNA using EpiMark ® Hot Start *Taq* Polymerase and NEBNext ® Multiple Oligos for Illumina ® (96 Unique Dual Index Primer Pairs) with: 30 s at 95°C; followed by 13 cycles of 15 s at 95 °C, 30 s at 61 °C, and 30 s at 68 °C. Purified PCR products were sequenced by the Centre for Genomic Research (University of Liverpool). Paired-end sequencing with 150 bp read length was conducted on a HiSeq 4000 sequencer (Illumina).

RRBS libraries were processed with Bismark (Krueger and Andrews, 2011) to obtain methylated and unmethylated counts. *G. aculateus* assembly version 5, ‘--bowtie2’, ‘--un’, ‘--ambiguous’, ‘-N 1’, ‘--no_dovetail’ options were used for the alignment step; ‘--paired-end’‘--no_overlap’, ‘--ignore 8’, ‘--ignore_3prime 3’, ‘--ignore_r2 10’, ‘--ignore_3prime_r2 15’ for the methylation extraction step. Samples with less than 100,000 reads were excluded, as were CpGs present in fewer than 16 individuals. For all remaining CpGs, M-values were calculated from raw count data using the formula:

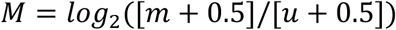

Where *m* is the count value of methylated reads and *u* is the count for unmethylated reads. CpGs on chromosome 19 -the stickleback sex chromosome -were removed to avoid sex differences in methylation state.

CpGs differentially methylated based on temperature were identified by redundancy analysis (RDA) using the *rda* function in the R package “vegan” (Dixon, 2003), using the model:

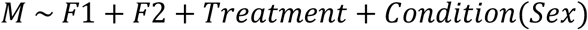

Where ‘Treatment’ is the combined F_1_ and F_2_ temperature (e.g. 12_12) and sex was used as a cofactor.

Outliers were identified as loci with scores exceeding ±2 standard deviations from the mean on either of the first two canonical axes. This cut-off was chosen by visual inspection of the screeplot of axis eigenvalues. Differentially methylated cytosines (DMCs) were identified as outlying CpGs that correlated significantly with F_1_ or F_2_ temperature. For both variables we identified DMCs by looping a biserial correlation analysis using the *cor.test* function in the “stats” R package (R Core Team, 2023) for each outlier, taking forward loci with p-values below 0.05.

### Identification of methylation-sensitive loci

We next quantified the extent to which methylation at specific DMCs modulated the relationship between genotype and phenotype. To ensure our analyses were computationally feasible, we reduced our set of DMCs to include those residing within a candidate region which had reads in at least 75 individuals (n=13,228). For each trait-model combination we then ran *scanone* QTL models with and without an individual DMC as a cofactor, looping this iteratively through all DMCs of interest. We calculated the difference in LOD score at each map marker between models with and without the DMC cofactor as in Broman and Sen (2009). Loci showing a LOD increase or decrease of at least 1 were classed as ‘methylation-sensitive loci’ (MSL) and were taken forward for further analysis. While we recognise that this threshold is somewhat arbitrary, a shift of LOD by 1 would represent a ten-fold change in log likelihood. Results using alternative thresholds are provided below. Genes containing a MSL (n=1533) were identified by aligning marker position to the V5 reference genome (Nath et al., 2021). Ensembl IDs were extracted and used to run GO analysis as before.

### Testing distribution and selection hypotheses

The role that methylation plays in driving TGP and WGP was quantified using a sliding window approach. The number of DMCs within each 10kb window (step size = 1kb) was measured using the *winscan* function in the “windowScanR” package (Tavares, 2025). We then tested if windows containing a QTL peak contained significantly more DMCs than those without a peak using a Wilcoxon Rank sum test, performed using the *wilcox.test* function in the “stats” R package (R core team, 2023). This was first performed on all QTL (n=51), followed by iterative runs on sets of QTL identified for unaltered morphology, TGP, and WGP respectively. We then repeated this approach tallying MSLs to test whether methylation-sensitivity was more prevalent in windows containing a QTL peak.

We then tested for evidence of selection driving plasticity, differential methylation, and methylation-sensitivity. Our sliding window approach was repeated to compare the mean F_st_ value in windows that did and did not contain a QTL peak, as above. ANOVAs were then run to test the correlation between mean F_st_ and the number of MSL and DMCs in each window. Finally, we repeated sliding window analysis on QTL identified for one of the four multi-generational treatment groups. This tested whether windows containing an ‘environmental QTL’ were enriched for signatures of selection, DMCs or MSL as before.

### Genetic architecture of methylation variation

To determine the genetic basis of genome-wide methylation variation we compressed DMC variation into a single variable amenable for QTL analysis. Separate runs of linear discriminant analysis (LDA) were performed for F_1_ and F_2_ DMCs respectively, using the *lda* function in the R package “MASS” (Ripley, 2011). This identified multivariate axes maximising variation between treatments at 12°C and 18°C for each generation to obtain canonical root scores to use as traits for QTL analysis. As a final step, variation in global methylation patterns was quantified, to allow comparisons with TGP-and WGP-associated methylation and provide a measure similar to morphology models. Principal component analysis (PCA) was run on the initial set of M-values, after regressing out the effects of sex with a linear model and taking each individual’s value for PC1 (explaining 11.22% of variation) as a trait. Inspection of eigenvalues (Supplementary Figure 1) showed a large drop in proportion of variance explained between PC1 (11.22%) and PC2 (3.7%), allowing us to take forward PC1 only. QTL analysis was run as before, with the exception that sex was not included as a cofactor having already been controlled for in the RDA model. Candidate genes, regions, and underlying GO terms were identified as above for morphological traits. Overlap between methylation QTL and QTL for TGP, WGP, and unaltered morphology were also tabulated as above.

## Results

### TGP and WGP have significant effects on morphology

Our *procD.lm* models found significant effects on morphological variation for TGP, WGP, and their interaction. WGP explained the greatest proportion of variation in morphology (13.02%, p=0.001) followed by TGP (1.59%, p=0.003) and the TGP*WGP interaction (1.46%, p=0.002).

### An ultra-dense linkage map for Icelandic threespine stickleback

Genetic map construction with Lep-Map3 produced the first ultra-dense linkage map for the SKR Icelandic stickleback populations (Supplementary Figure 2). Our final map was 2981.99 centimorgans (cM) in length and contained 11,342 markers genotyped with 99.7% coverage in 294 individuals (Supplementary Table 2).

### TGP and WGP involve distinct genes and biological functions

TGP and WGP involved distinct genetic regions, potentially allowing for the two forms of plasticity to evolve independently. Partial pairwise correlation scores between TGP and WGP models for the same traits showed weak and often negative correlations, with an average correlation of 0.088 (Figure 2A; for raw values see Supplementary Table 3). This was lower than relationships between morphology and TGP (0.346) and WGP (0.386) respectively.

**Figure 2.**
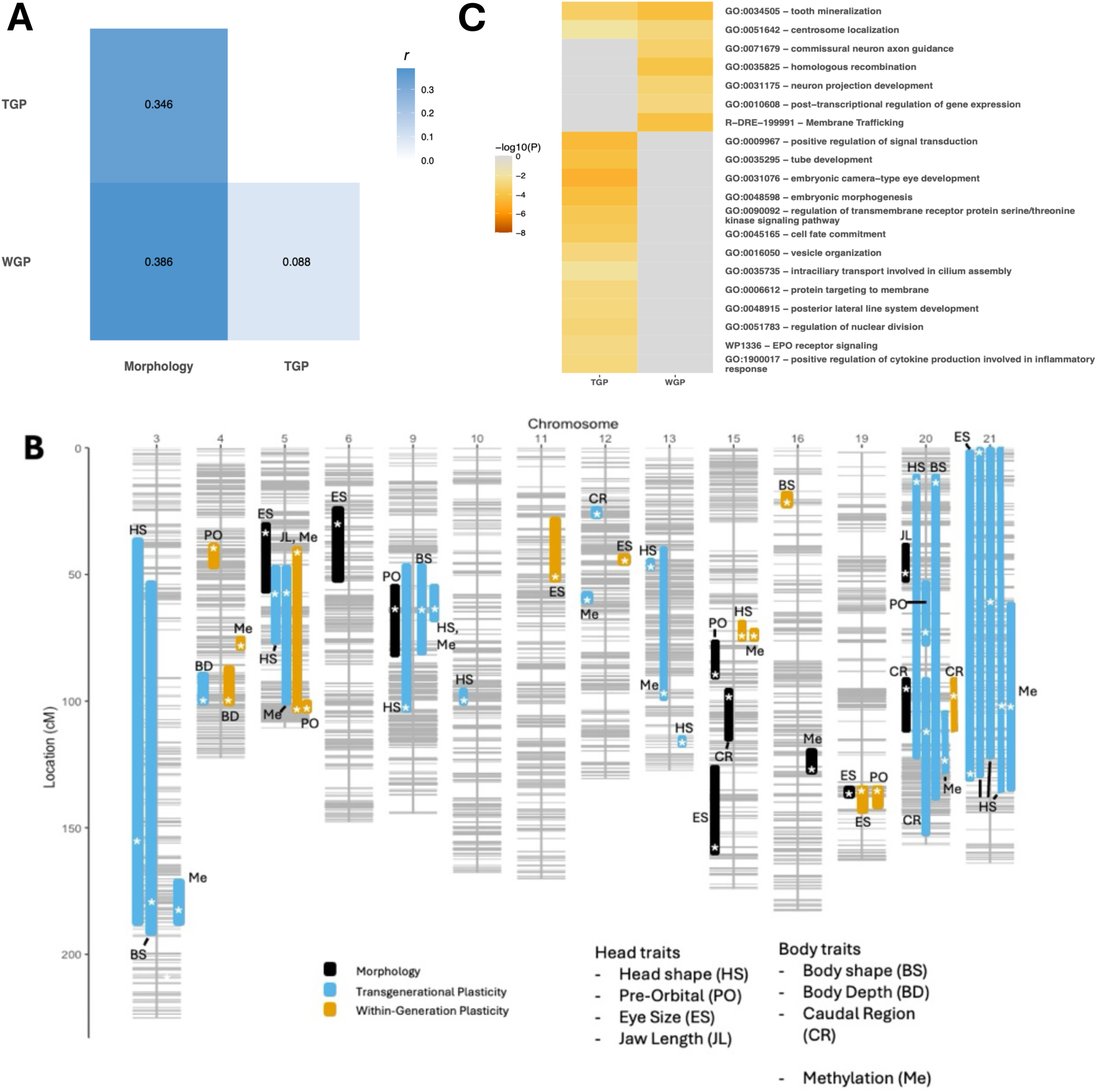
Transgenerational plasticity (TGP) and within-generational plasticity (WGP) are associated with distinct genes and biological functions. **A)** TGP and WGP are weakly correlated at the genetic level. Pairwise partial correlation scores comparing LOD scores obtained from TGP and WGP models showed an average correlation (*r*) of 0.088 across eight morphology and methylation-based QTL models. Models of morphology and TGP and WGP showed stronger correlations. Strength of correlation between pairwise model comparisons is represented by shading (darker = stronger correlation). **B)** Distribution of QTL across the genetic map. 1.5-lod confidence intervals around QTL are plotted in centimorgan (cM). Intervals are colour-coded based on model type (black – morphology; blue-TGP; orange – WGP). QTL peaks are depicted with asterisks. Only chromosomes containing at least one QTL are depicted. **C)** TGP and WGP involve distinct biological functions. Terms enriched in the set of all genes falling within candidate regions of TGP and WGP QTL are enriched in one form of plasticity alone. Gene ontology (GO) analysis was performed in metascape using *Danio rerio* gene IDs. Hue represents the –log10 p-value associated with each term, with darker colours representing lower p-values and hence greater statistical power. The top 20 enriched terms only are shown.

Furthermore, TGP and WGP QTL were differentially distributed across chromosomes. Partial pairwise correlation scores for the chromosomal distribution of TGP and WGP QTL showed a negative correlation, but this was small and not significantly different from zero (r = -0.014, p=0.95).

Of 36 QTL for TGP or WGP, on only three occasions did TGP and WGP map to the same chromosome for the same trait (Figure 2B). Of the 18 pairwise combinations of TGP and WGP QTL falling on the same chromosome, confidence intervals overlapped in 12 cases, although most of these overlaps were driven by large confidence intervals on chromosomes 5 and 20, with QTL peaks for TGP and WGP only falling within 2.5Mb on two occasions (Supplementary Table 4).

Beyond their largely exclusive genomic locations, TGP and WGP were enriched for different biological processes, supporting a functional delineation. Gene ontology analysis performed on all genes within candidate regions for TGP or WGP QTL showed that, of 135 statistically significantly enriched GO terms (logP ≤ -2), only 7 (5%) were significantly enriched in both gene sets (Supplementary Table 5). Metascape’s algorithm for dealing with redundancy -clustering terms based on gene membership and taking the term with the lowest p-value as representative (Zhou et al., 2019) -finds that only two of the top 20 best-supported terms are enriched for both TGP and WGP (Figure 2C), and only four of the top 100. Confidence intervals around TGP QTL were enriched in GO terms for morphogenetic processes (GO:0048598: embryonic morphogenesis; GO:0031076: embryonic camera-type eye development) and developmental regulation (GO:0009967: positive regulation of signal transduction; GO:0140966: piRNA-mediated heterochromatin formation) (Supplementary Figure 3a). Conversely, WGP genes were associated with metabolism (GO:0140053: mitochondrial gene expression; GO:0055088: lipid homeostasis) and gene expression regulation (GO: 0010608: post-transcriptional regulation of gene expression; dre03083: polycomb repressive complex) (Supplementary Figure 3b).

### Underlying genes and biological functions are influenced by parent and offspring temperature

The genetic architecture of morphological traits was influenced by the cumulative effects of parental and offspring temperatures. Partial correlation scores comparing the four treatment groups yielded values around zero (Figure 4; Supplementary Table 6). This was true of all treatment-treatment comparisons, with values not being increased by a shared F_1_ or F_2_ temperature. GO analysis showed that of 106 statistically significantly enriched terms, 93% (99) were significantly enriched (logP ≤ -2) in one of the four treatment groups only. No GO term was significantly enriched across three or more treatment groups. Only one term was identified as non-redundant by the metascape algorithm - GO:0042451: purine nucleotide biosynthetic process - was shared across multiple treatment groups. Functions enriched for the 18_18 treatment group involved signalling (GO:0023056: positive regulation of signalling; wp341: Nodal signalling pathway) and metabolism (R-DRE-9711097: cellular response to starvation; dre00190: oxidative phosphorylation). Meanwhile, terms enriched at 12_12 were principally involved in stress responses (GO:0001666: response to hypoxia; GO:0034599: cellular response to oxidative stress).

**Figure 3.**
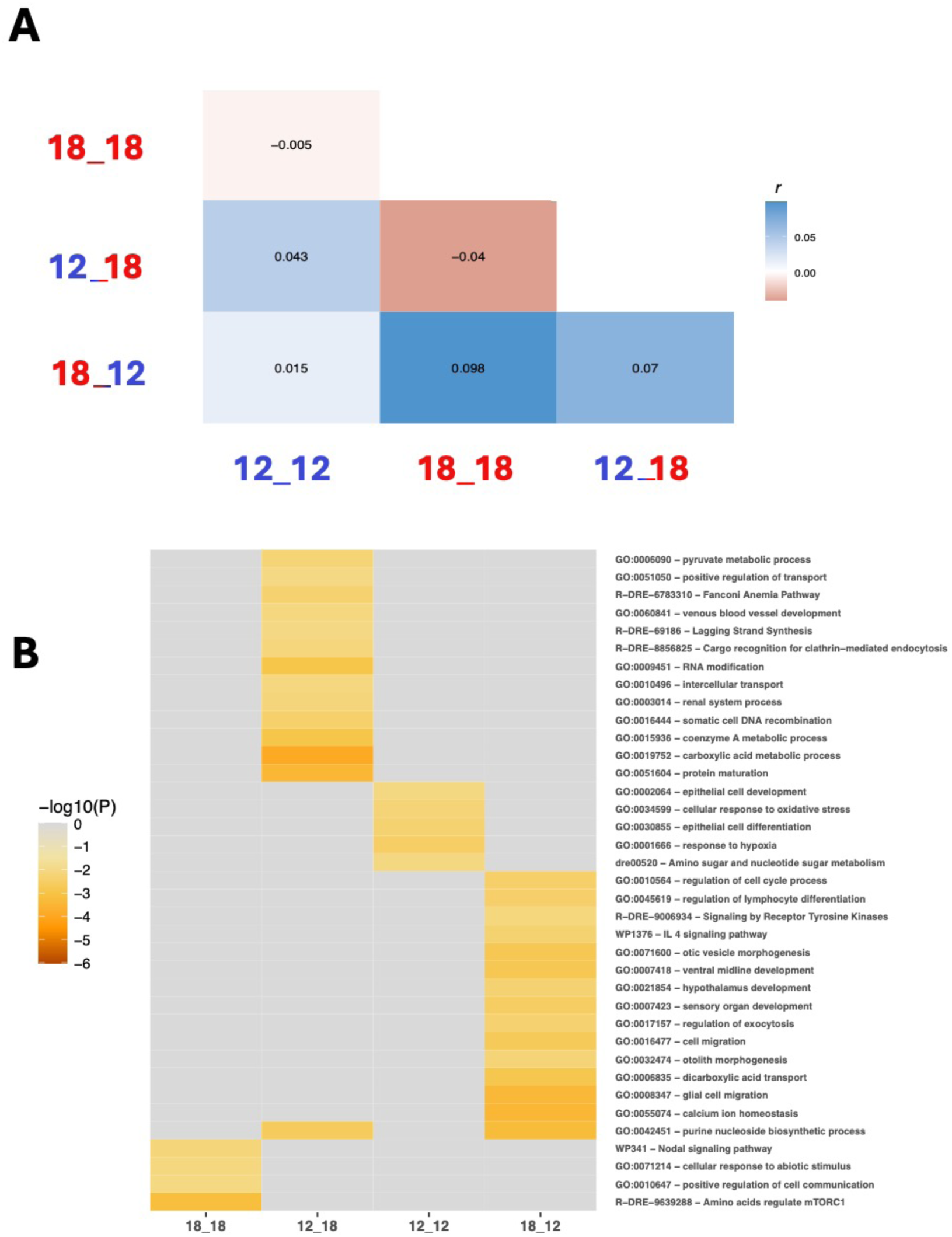
WGP and TGP shape the genetic architecture and biological functions underlying morphology. **A)** Pairwise partial correlation models for our 4 multi-generational treatment groups showed weak and often negatively correlations (*r*) for LOD scores across eight morphology and methylation-based traits. Treatment groups take the format ‘12_12’ where the first number represents the F1 temperature while the second number represents the F2 temperature. Generations spent at 12°C and 18°C are represented in blue and red respectively. Grid squares are shaded based on the sign and magnitude of correlation, with squares tending towards blue and red representing positive and negative correlations respectively. **B)** Enriched biological functions vary between treatment groups as indicated by gene ontology analysis. This includes all genes falling within confidence intervals for QTL associated with any of the four treatment groups with all non-redundant enriched terms (logP ≤ -2) are depicted. Hue represents the –log10 p-value associated with each term, with darker colours representing lower p-values.

**Figure 4.**
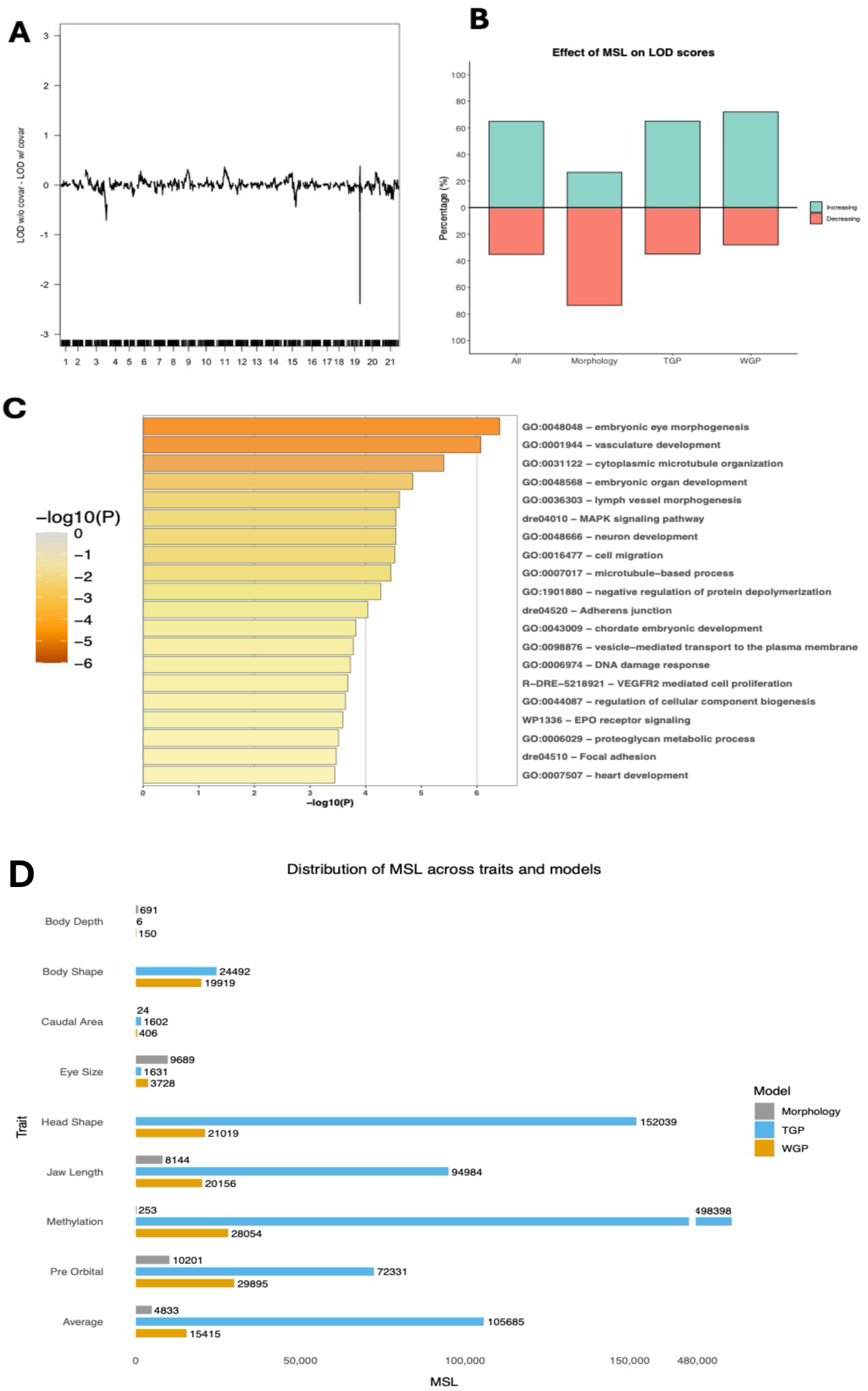
Methylation alters the relationship between genotype and phenotype. **A)** Demonstration of the Methylation-sensitive loci (MSL) concept. MSL are defined as genetic map markers that change LOD score by one in either direction with the addition of a differentially methylated cytosine (DMC) as a cofactor. In one such case, addition of a DMC cofactor reduced the LOD score by more than 2 points at one locus compared to a comparative model without the cofactor. The QTL ‘valley’ on chromosome 21 corresponds to the gene *tle3a*, associated with wnt signalling, cold-induced thermogenesis, and interactions with histone methyltransferases. **B)** Adding a DMC cofactor tended to increase LOD scores at methylation-sensitive markers. This was true across all QTL, those associated with TGP and WGP, but the opposite trend was found for QTL associated with morphology **C)** Genes containing MSL were implicated in morphogenesis and developmental signalling. GO analysis was performed in metascape using all genes found to contain a methylation-sensitive locus. The top 20 enriched terms only are shown, with hue representing –log10 p-values. **D)** The number of MSL varies by trait and model type. The number of MSL is shown on the for each trait and model combination, with averages plotted at the bottom. Models are colour-coded (Grey – Morphology; Blue – TGP; Orange – WGP).

QTL associated with any of the four multi-generational treatments showed signatures of selection. 10kb windows containing a QTL peak had marginally, but significantly higher mean F_st_ scores than windows that did not (0.0249 vs 0.0237, p=6.3×10^-5^). These windows also contained significantly more DMCs (16.3902 vs 7.6661, p=2.2×10^-16^) and MSL (1.5125 vs 0.1959, p=2.2×10^-16^) than windows that did not contain a peak.

Unaltered morphology QTL tended to show different inheritance patterns across treatment groups (Supplementary Figure 4), demonstrating that the environment determines what heritable variation is exposed to selection through the phenotype. Of the ten QTL for unaltered morphology found previously, only two showed consistent allele sign effects across all four treatment groups. On four occasions, HSE values had opposite signs in 12_12 and 18_18 individuals. Several QTL, such as one for caudal region size on chromosome 20, have large allele effects in one treatment group only and modest effects in all others, highlighting the capacity of genetic variation to be ‘cryptic’ under certain environmental conditions.

### Genotype-phenotype correlations are modulated by methylation

With individual DMCs incorporated into QTL models as cofactors we demonstrated that methylation had pervasive effects on genotype/phenotype relationships, including their plasticity. Using a subset of 13,228 DMCs, we found 997,812 marker-DMC-trait combinations that changed LOD scores by at least 1 in either direction (Figure 4A). Such effects were extensive with 8034 markers on our genetic map (70.83%) found to be methylation-sensitive based on these criteria, including all QTL peaks. The addition of a DMC as a cofactor tended to raise LOD scores (64.79% increased, 35.21% decreased), suggesting their addition provided increased explanatory power. This was true when looking at models for TGP (65.09 vs 34.94%) and WGP (71.96 vs 28.04%), but unaltered morphology models showed the opposite trend (26.53 vs 73.47%) (Figure 4B). Genes containing a MSL were involved in embryonic eye morphogenesis (GO:0048048), cell migration (GO:0016477), DNA damage response (GO:0030201) and the MAPK signalling pathway (dre04010) amongst other terms (Figure 4C). Genes containing large-effect MSL include *tle3a*, know to regulate cold-induced thermogenesis (Villanueva et al., 2013) and interact with histone methyltransferases (Kalita et al., 2024).

TGP models contained several-fold more MSL on average than WGP or unaltered morphology models (generalised linear model, *Z*=1749.8, *p*<2.2×10^-16^), although this varied by trait (generalised linear model, z=196.21, p<2.2×10^-16^) (Figure 4D). MSL density also related positively to F_st_ scores within 10kb windows (*t*=152.98, *p*<2.2×10^-16^), suggesting selective pressures acting on methylation-sensitive genes. Moreover, 10kb windows containing a QTL contained more MSL (1.8147) than windows that did not (0.1962) (*p*<2.2 x10^-16^), although we used only DMCs falling within candidate regions, which likely imposed a bias (Table 1). While this applied for sets of QTL associated with each model, windows containing TGP QTL indicated a more prominent role for methylation by containing significantly more MSL (2.0044) than windows containing WGP QTL (1.6250) (*p*<2.2×10^-16^)

**Table 1.** Sliding window analysis comparing 10kb windows with (left) and without QTL (right). Mean values are shown, with asterisks denoting statistical significance: ns p >0.05; * *p* < 0.05; ** *p* < 0.01; *** *p* < 0.001. Fst = Mean Fst score in a genomic window; DMCs = Mean number of differentially methylated cytosines; MSL = Mean number of methylation-sensitive loci. TGP = Transgenerational plasticity, WGP = Within-generation plasticity. Environmental QTL are associated with one of the four multi-generational treatment groups.

| **Model** | **F<sub>st</sub>** | **DMCs** | **MSL** |
|---|---|---|---|
| **Combined** | 0.0279<sup>***</sup> - 0.0237 | 10.1572<sup>***</sup> - 7.6737 | 1.8147<sup>***</sup> - 0.1962 |
| **Morphology** | 0.0242<sup>ns</sup> - 0.0238 | 8.4778<sup>***</sup> - 7.6758 | 1.5600<sup>***</sup> - 0.1975 |
| **TGP** | 0.0318<sup>***</sup> - 0.0237 | 12.6769<sup>***</sup> - 7.6734 | 2.0044<sup>***</sup> - 0.1968 |
| **WGP** | 0.0289<sup>*</sup> - 0.0238 | 5.7455<sup>*</sup> - 7.6764 | 1.6250<sup>***</sup> - 0.1975 |
| **TGP vs WGP** | 0.0318<sup>ns</sup> - 0.0289 | 12.6769***- 5.7455 | 2.044<sup>***</sup> - 1.6250 |
| **Environmental QTL** | 0.0249<sup>***</sup> - 0.0237 | 16.3902<sup>**</sup> - 7.6661 | 1.5124<sup>***</sup> - 0.1959 |

### QTL show evidence of selection and differential methylation

Sliding window analysis provided evidence for selection on TGP, highlighting a role for methylation in guiding adaptation. This included 10kb windows with QTL peaks having more CpGs (mean = 10.1557) that were differentially methylated based on temperature than windows without (mean = 7.6737) (Mann-Whitney; *p*=5.5059×10^-16^) (Table 1). This was similarly true for windows containing QTL associated with unaltered morphology (8.4778; *p*=5.458×10^-4^) and TGP (12.6769; *p*=4.705×10^-14^) while windows containing WGP QTL contained significantly fewer CpGs than windows without them (5.745; *p*=0.01268). Overall, windows containing a QTL had a higher mean F_st_ value (0.0279) than windows that did not (0.0242) (*p*=6.399×10^-5^). However, this was only true for windows containing QTL associated with TGP (0.0318; *p*=1.22×10^-5^) or WGP (0.0289; *p*=0.03409).

### Methylation genes are involved in plasticity and metabolism

We sought to map the genetic basis of methylation to understand the role it plays in morphological plasticity. The average read mapping efficiency for the Bismarck alignment was 56.52, with an average percentage of CpG methylation of 61.4%. Mapping the genetic basis of methylation identified genes associated with craniofacial development (GO:0034505: tooth mineralisation; GO:0031076: embryonic camera-type eye development), chromatin organisation (GO:2001251: negative regulation of chromatin organisation; GO:0140966: piRNA-mediated heterochromatin organisation) and metabolism (R-DRE-2142789: Ubiquinol biosynthesis; GO:0008654: phospholipid biosynthetic process). Of 138 terms surpassing significance (logP ≤ -2), all but four (2.89%) are distinct to either TGP or WGP methylation, of which three are non-redundant as per the metascape clustering algorithm (Supplementary Figure 5).

Confidence intervals for methylation and morphological plasticity QTL overlapped frequently, suggesting a shared genetic component. Overlaps were observed on 20 of 30 occasions where models for methylation and either TGP or WGP fell on the same chromosome, with these overlaps typically occurring when the generation (e.g. TGP/WGP) matched across model types. In four instances QTL peaks were implicated in both morphological plasticity and methylation (2 for TGP, 2 WGP), with generations matching in all cases. Three of these QTL were identified as candidates for multiple traits in the craniofacial region, suggesting pleiotropy. On five further occasions peaks fell within 1.5Mb, with generations matching in all but one instance. Regions for TGP and WGP methylation mapped to different chromosomes with the exception of chromosome 15, where overlapping intervals were observed and respective peaks fell within 1Mb.

## Discussion

We investigated the genetic architecture of TGP and WGP in response to temperature, incorporating methylation variation to understand how these information channels interact and evolve in populations undergoing thermal adaptation. We found that TGP and WGP were associated with distinct regions of the genome, and with genes conferring distinct biological functions. Our full factorial design demonstrated that trait genetic architectures were dependent upon parental and offspring environments, their interaction, and the methylome.

Finally, our results suggest that methylation drives TGP, is targeted by selection and modulates genotype-phenotype relationships. Overall, our findings highlight the multiple streams of information that determine the genetic basis of phenotypic variation, and their interactions, which themselves can be targeted by selection in the process of adaptive divergence.

### TGP and WGP represent distinct evolutionary processes

The distinct genetic architectures of TGP and WGP provide a capacity for the two forms of plasticity to evolve independently, limiting potential constraints from cues arising within and across generations. Pairwise correlation scores between TGP and WGP LOD values for the same trait were weakly or negatively correlated, with QTL mapping to distinct sets of chromosomes and confidence intervals rarely overlapping. Theory suggests that TGP and WGP will be adaptive or maladaptive in different scenarios, typically based on the correlation between parent and offspring environments (Colicchio and Herman, 2020). Modelling approaches tend to find that selection on TGP and WGP is negatively correlated in most cases (Clement et al., 2023). The decoupling of TGP and WGP at the genetic level in our data may allow populations to find the optimal balance between both forms of plasticity.

Accordingly, Díaz and Matzkin (2025) found that genes associated with TGP and WGP had opposing transcriptional trajectories; genes upregulated in response to the parental environment were downregulated in response to the offspring environment, and vice versa. QTL (Che et al., 2024) and gene expression studies (Déjeux et al., 2025; Hales et al., 2017) have similarly identified distinct sets of genes associated with each form of plasticity. Thus, the genetic decoupling of TGP and WGP may allow populations to adapt to complex and fluctuating environments.

TGP and WGP were associated with distinct biological processes, suggesting the targets of selection vary according to the generation in which a cue is experienced. Indeed, 95% of enriched GO terms were unique to one form of plasticity, supporting theory suggesting that TGP and WGP are involved in different processes and mechanisms (Bell and Hellmann, 2019; Jablonka and Raz, 2009) and in line with previous QTL data from plants (Che et al., 2024), epigenetics (Ryu et al., 2020) and gene expression (Hales et al., 2017). TGP was associated with the morphogenesis of anatomical features, while WGP was implicated in metabolism and gene expression regulation. Related to this, recent insights from Icelandic stickleback show that patterns of covariation are dominated by TGP (Brachmann et al. in prep), perhaps because trait integration is determined early in ontogeny when transgenerational information may be more influential. In support, TGP bears a relatively stronger effect on traits expressed early in life in plants (Wadgymar et al., 2018) and influences morphological rather than behavioural traits in response to predator exposure in insects (Sturiale and Bailey, 2023). Thus, we suggest that TGP is providing a morphological ‘scaffold’ which is modified and buffered by WGP. This view frames the two forms of plasticity as distinct processes involving different genes and biological functions, representing different evolutionary targets.

### Cryptic genetic variation drives the evolution of thermal plasticity

How the environment can determine what allelic variation is exposed to selection is a central question for both evolutionary-developmental biology (evo-devo), and increasingly conservation biology. We found that the genetic basis, and biological functions underlying trait variation, were highly dependent on the thermal environment of both the parent and offspring. Indeed, correlation scores between the four treatment groups were weak or negative, with 92.5% of enriched GO terms identified for one treatment group only. Evidence for environmental specificity of genetic architectures has been provided previously, with between 33 and 83% of QTL showing QTL-by-environment interactions (Lafuente and Beldade, 2019). By using multi-generational treatments, we found that what genetic information is drawn upon depends on both the parental and offspring environments, and their interactions. Empirical studies frequently show non-additive interactions between TGP and WGP (Neylan et al., 2025) suggesting molecular mechanisms underpinning cue integration. At the genetic level, Che et al. (2024) demonstrated that the genetic architecture of TGP depended on the offspring environment, and vice-versa, highlighting the interactivity between these two streams of information. Overall, low correlation scores between model types and treatment groups point towards thermal plasticity arising through highly polygenic and environmentally sensitive genetic architectures, leading us towards a view of genes as ‘potential information’ (Danchin, 2019), drawn upon depending on the environment.

If plastic responses are driven by genes that have phenotypic effects in some environments but not others, this ‘cryptic’ genetic variation (CGV) (Gibson and Dworkin, 2004) will create challenges for those applying approaches relying on a fixed relationship between genotype and phenotype, such as genetic ‘offset’ models (Ahrens et al., 2025; Lind and Lotterhos, 2024). For instance, our data implicated genetic variants at 12-degrees that were phenotypically silent at 18-degrees, and vice-versa. Thus, loci conferring an adaptive fit to current conditions may not be relevant in a warmer world, while those that will drive adaptation may be invisible in current environments. Inheritance patterns of morphology QTL were also found to vary across the four treatment groups, highlighting the capacity of the environment, at the most fundamental level, to change how genes are exposed to selection. Indeed, genomic windows containing these ‘environmental QTL’ had significantly higher F_st_ scores than windows that did not, suggesting exposure to selection. Understanding these dynamics and viewing organisms as developmental systems, rather than tabulations of allele frequencies, will be central to predicting the speed and accuracy of climate change responses.

### Methylation is targeted by selection and underlies TGP

Integrating genetic and epigenetic data revealed that methylation drives TGP and occurs in regions of the genome facing relatively stronger selection. QTL and gene ontology terms for morphological and methylation plasticity frequently overlapped, supporting a mechanistic role, and highlighting that loci underlying methylation can be viewed as candidates for TGP. Genomic windows associated with plasticity, particularly TGP, had significantly more differentially methylated cytosines (DMCs) and methylation-sensitive loci (MSL), demonstrating this mechanistic link further. In addition, genomic windows associated with environment QTL had notably more DMCs than null expectations, further suggesting a role for methylation in exposing the genome to environmental variation. The generality of this finding, with the same locations appearing in independent analyses, validates our experimental approach. We combined a novel morphometric technique, extract TGP and WGP, with a more established approach for identifying methylation variation in response to explanatory factors (redundancy analysis) and found both approaches independently led us to the *same* genomic regions. Thus, by combining two sources of data, we have been able to verify the genetic and epigenetic mechanisms of plasticity and document their interactions.

TGP, WGP, and methylation were enriched for signatures of selection in wild populations, suggesting that these processes are evolving in response to warming. Genomic windows containing plasticity QTL showed significantly higher mean F_st_ scores than windows that did not, and this was especially true for TGP. F_st_ scores were also positively correlated with the number of DMCs and MSL within windows, the counts of which are both higher in TGP windows. This suggests methylation and TGP are under selection and implicated in processes underlying thermal plasticity. Furthermore, these results suggest that the findings from our hybrid cross, seeded by two individuals, are applicable to wild populations. Our results contradict theory that phenotypic plasticity hinders evolution by shielding pertinent variation to selection (Sommer, 2020; Williams, 1966). Rather, plasticity and methylation represent targets for selection to act on, driving the evolution of plastic responses upon environmental change.

Combining genetic and epigenetic data into the same QTL models demonstrates that methylation drives plasticity by modulating genotype-phenotype relationships. Over 70% of mapped markers, including all QTL peaks, were found to be ‘methylation-sensitive’, meaning their effect on morphological phenotypes were modulated by methylation. The addition of DMCs as cofactors tended to increase LOD scores suggesting that most QTL studies could be underestimating the relationship between genotype and phenotype, while methylation itself could be increasing the efficiency of selection to act on genetic variation. MSL were enriched in biological functions implicated in morphological plasticity. This included eye development, a highly plastic divergent trait in geothermal populations (Pilakouta et al., 2023; Smith et al., 2024), and MAPK signalling, identified as divergent in wild populations (Brachmann et al., 2026). Finally, the number of MSL in a genomic window was positively correlated with F_st_ scores, suggesting that methylation-sensitivity is under selection directly, or is involved in processes or regions which are.

TGP, driven by methylation, may be favoured for several reasons. Firstly, several QTL for methylation were also implicated in the plasticity of numerous morphological traits, suggesting a role for methylation in driving co-ordinated plastic changes (Navon et al., 2021; West-Eberhard, 1989). In the context of wild geothermal stickleback, temperatures are fairly stable from one generation to the next. Thus, parental and offspring environments are ‘autocorrelated’, with inherited information likely to be relevant, a scenario in which the evolution of TGP is supported (Clement et al., 2023). Furthermore, if transgenerational cues persist for two or more generations, this will allow more opportunity for selection to act and refine plasticity. Finally, epigenetic marks and F_st_ scores may be correlated due to the relationship between methylation and germline mutations. Methylation can act as a precursor of genetic change as methylated cytosines are approximately 12 times more likely to mutate than unmethylated cytosines (Guerrero-Bosagna, 2020). This has led to theories that methylation can act as a conduit between plastic and genetic change (Anastasiadi and Piferrer, 2019; Danchin et al., 2019; Ord et al., 2023) which our data supports. Hence, methylation hotspots may generate more *de novo* mutations which can be selected on, partly in response to environmental factors.

### A role for metabolism in morphological plasticity

Despite measuring variation in morphological plasticity, genes and terms relating to metabolism appeared throughout. Notably, we identified several metabolic processes here that have been highlighted in previous work on this system (Brachmann et al., 2026). These co-occurrences suggest a role of TGP and WGP in thermal adaptation, or at least shared mechanisms. Firstly, our results highlighted MAPK genes which are known to be differentially expressed in geothermal and ambient fish (Brachmann et al., 2026). The MAPK pathway is associated with cell proliferation and neural crest cell development (Dinsmore and Soriano, 2018) and has been implicated in other examples of adaptation to warmer temperatures (Campbell-Staton et al., 2021; Jacobs et al., 2024) making it a candidate pathway in the Icelandic stickleback system. In this study, MAPK-related GO terms were identified for WGP and methylation-sensitive loci, suggesting a potential role in morphological plasticity. Notably, we found the gene *ncoa2,* known to be regulated by the MAPK pathway (Burgermeister and Seger, 2007), directly under a QTL peak underlying TGP in both head shape and methylation. This gene, associated with appetite regulation and lipogenesis (Lu et al., 2015; Rosen, 2016), has previously been identified as a genomic outlier (Brachmann et al., 2026); *ncoa2* was one of several candidate genes previously found in a putative inversion on chromosome 21 (Brachmann et al., 2026). GO analysis of this inverted region found terms relating to metabolism, hypoxia, and cellular repair, all of which were identified as enriched in our analyses. Finally, we also identified terms relating to insulin and lipid metabolism, believed to aid adaptation to geothermal environments (Brachmann et al., 2026)

While the importance of metabolic change in a warmer world is evident, why such terms appear in analyses of morphological variation is less clear. One potential explanation is the action of pleiotropic loci influencing morphological and metabolic plasticity. Such loci may be targeted by selection for the ability to drive co-ordinated phenotypic shifts. The presence of loci implicated in morphological plasticity and methylation suggests a potential mechanism, where changes to methylation patterns having multi-dimensional phenotypic effects. Another explanation is that metabolism could be considered as part of development, especially in stressful environments. Development is an inherently metabolically demanding process (Confavreux et al., 2009; Kordas et al., 2023), thus ensuring physiological conditions may be an important, yet overlooked component of morphological plasticity. Temperature, as a strong and multi-dimensional stressor, may therefore favour genetic changes that lead to changes in both morphology and metabolism.

### Concluding remarks

Effective conservation practice requires an understanding of how phenotypes are produced and acted on in changing environments (Campbell et al., 2017; Feiner et al., 2021). Our results show that development integrates information from the genome, epigenome, and past and present thermal environments to produce the variation that drives evolution. This speaks to the environment’s role in generating phenotypic variation, while also determining what genetic variation is shielded or exposed to selection. Evidence for selection on both methylation and TGP, combined with weak or negative correlations across treatment groups, reaffirms that adaptive evolution acts on the phenotypic interface between the environment and development, rather than specific loci (Mayr, 1997). Thus, relationships between the genotype, environment, and phenotype may be targeted by evolution in the early stages of adaptation (Simpson, 1953; Sultan, 2015), particularly in stressful environments. Important interactions between TGP and WGP, methylation and genetic variation were also observed, highlighting the limitations of studying any such process in isolation. We would therefore recommend conservationists to integrate system-level approaches into practice, while moving away from approaches based on the action of individual loci. In sum, our results show the cycles of reciprocity between generative and selective forces that drive evolution in changing environments (Lala et al., 2024; Stansfield and Parsons, 2024).

## Acknowledgments

We would like to thank the animal technicians at the University of Glasgow including Ross Phillips, Alastair Kirk, and the late Graham Law for their help with animal husbandry and aquarium maintenance. We would also like to thank members of the Parsons lab for their constructive comments on earlier drafts of the manuscript, and Alex Brumwell for technical assistance in the lab. Pasi Rastas and Karl Broman provided helpful input on genetic map construction and QTL mapping respectively. We thank Kári Heiðarsson for his help with fieldwork logistics, and for access to Áshildarholtsvatn and for keen interest in the project we thank Sigurður Guðjónsson and Guðrún Kristmundsdóttir

## Ethics statement

Our study adheres to the ASAB/ABS Guidelines for the Use of Animals in Research, the institutional guidelines at the University of Glasgow, and the legal requirements of the UK Home Office (Project License P89482164)

## Supplementary figures

**Supplementary Figure 1.**
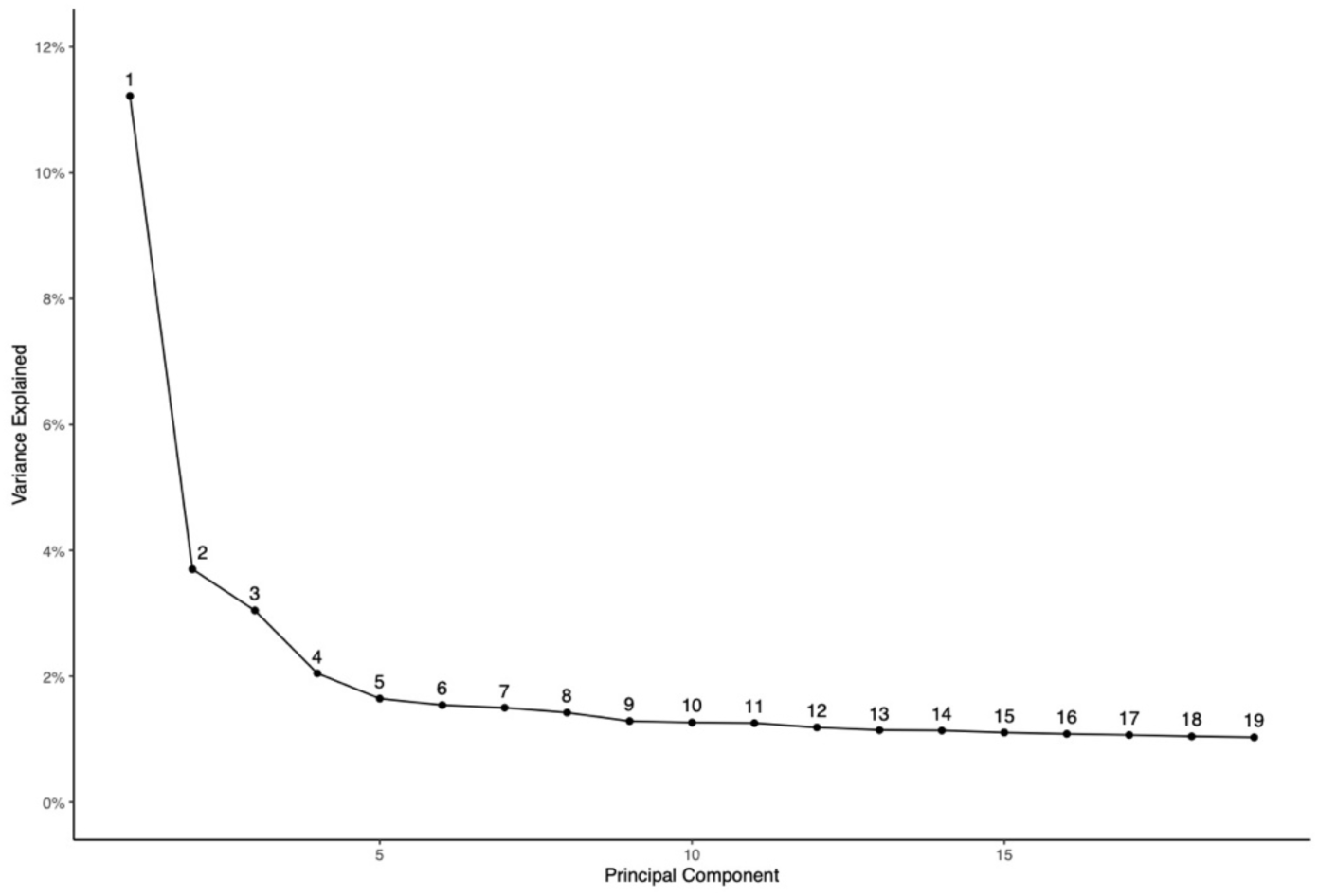
Methylation PC1 explained 11% of variation. Principal component analysis was run on raw m-values, with the percentage of variation explained by each visualized in a screeplot. PC1 was taken forward as a trait for QTL analysis. Only PCs explaining more than 1% of variation are plotted

**Supplementary Figure 2.**
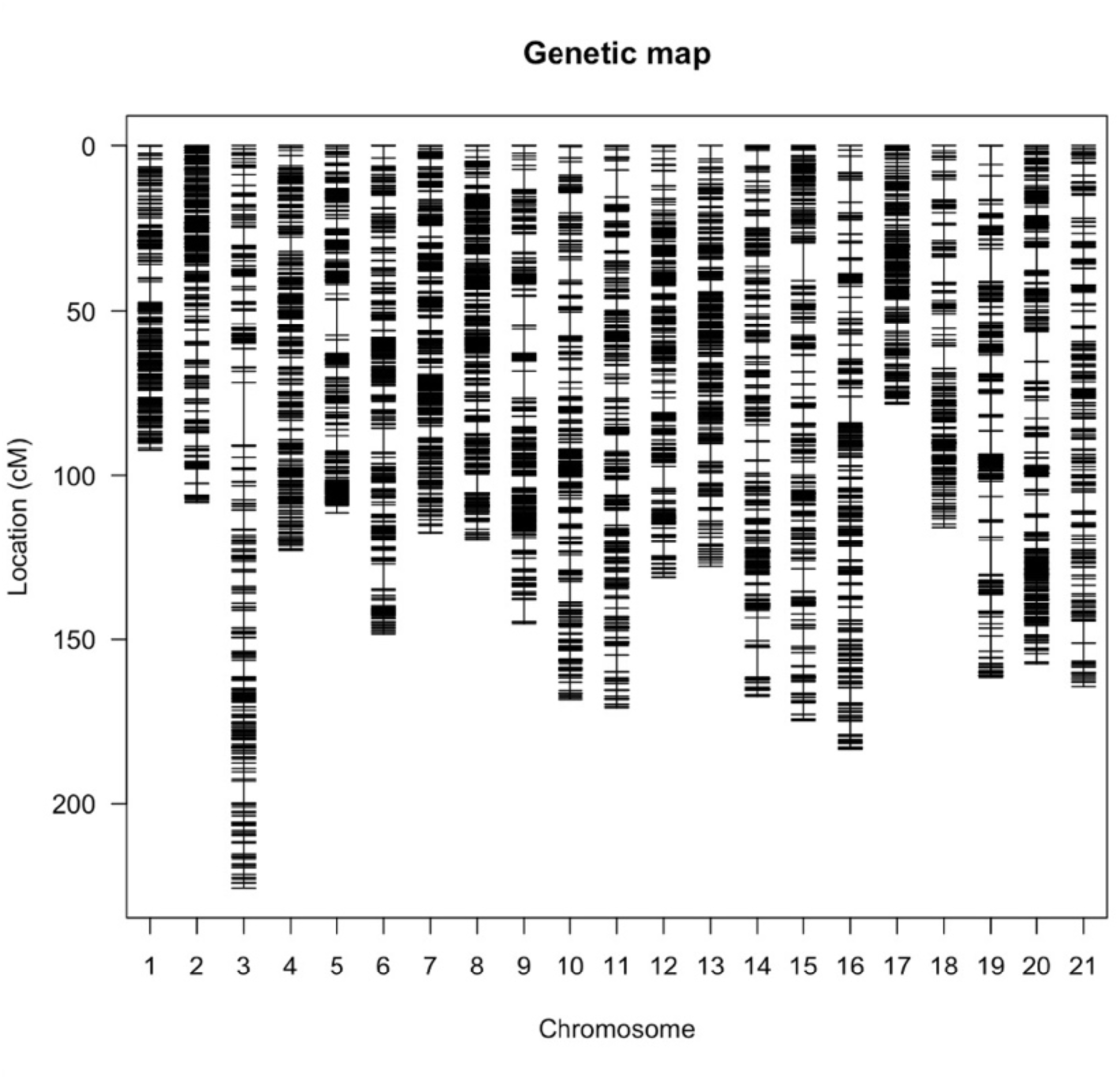
An ultra-dense genetic map for the Icelandic threespine stickleback. Genetic map construction was performed using Lep-Map3, from RRBS derived genotype data. Markers were ordered on each chromosome by physical position, aligned to the stickleback V5 reference genome. The final map contained 11,342 markers genotyped with 99.7% coverage in 294 individuals.

**Supplementary Figure 4.**
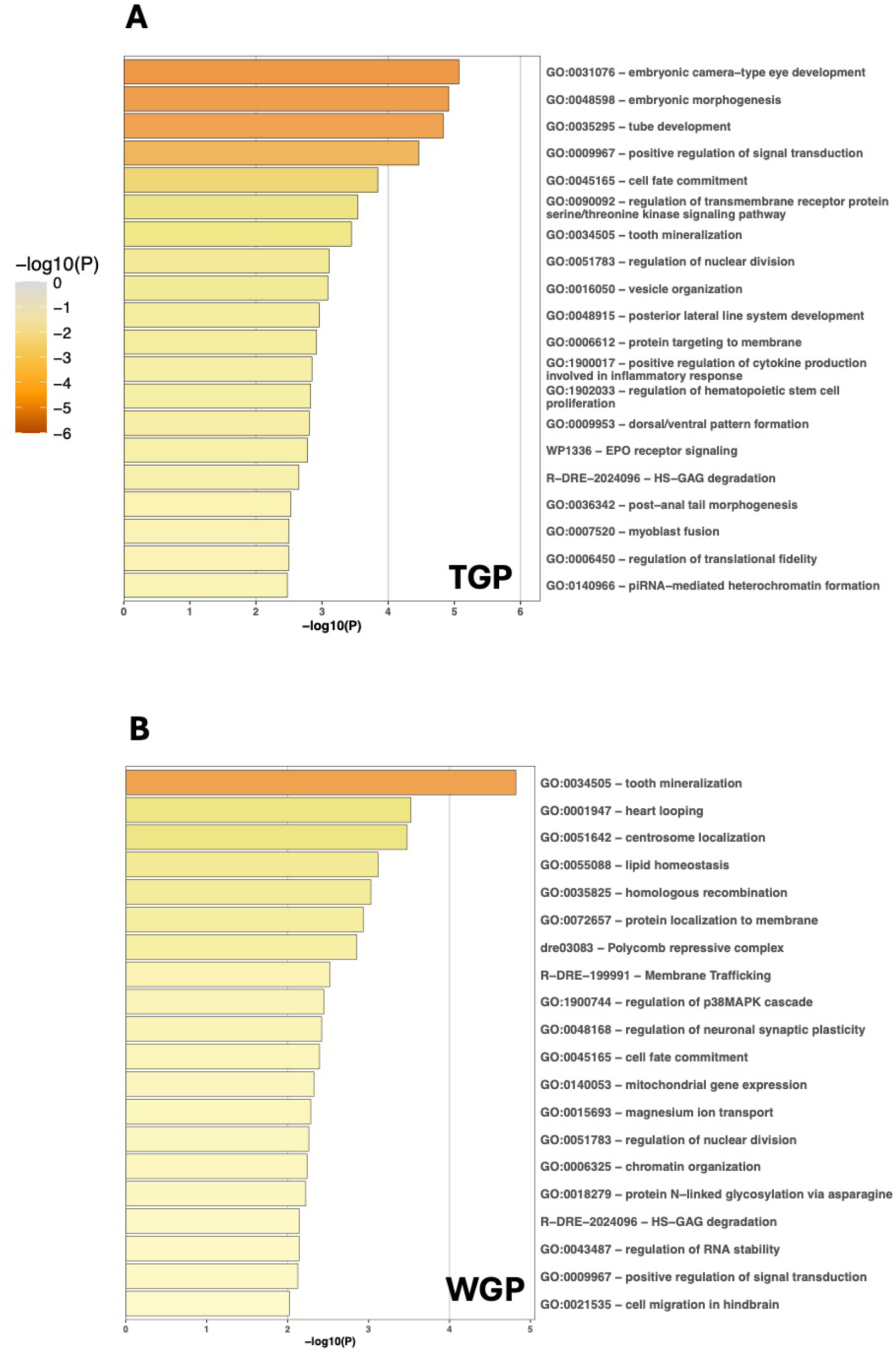
TGP and WGP are associated with distinct biological functions. Top 20 terms enriched in candidate regions associated with **A)** TGP and **B)** WGP. Gene lists were submitted separately and GO analysis was performed using metascape with *Danio rerio* gene IDs. Hue represents the -log10 p-value associated with each term, with darker colours representing lower p-values and hence greater statistical power.

**Supplementary Figure 5.**
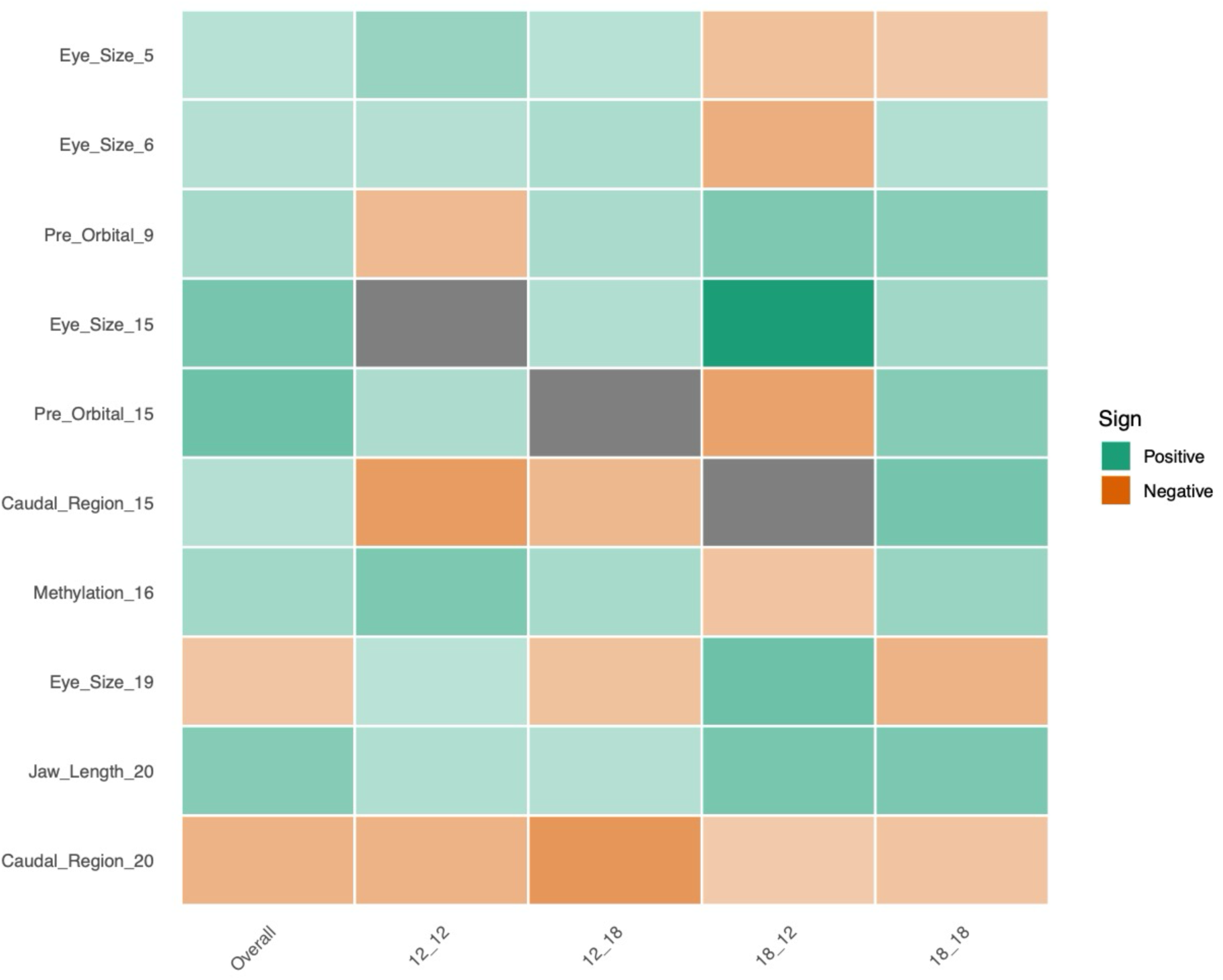
Inheritance patterns differ across multi-generational treatment groups. The homozygous substitution effect (Mean AA phenotype – Mean BB phenotype) is shown for each morphology QTL and for each multigenerational treatment group. Treatment groups were named in the format ‘12_12’ where the first number represents the F_1_ temperature while the second number represents the F_2_ temperature. Squares represent the sign and magnitude of the allele substitution effect, scaled against the standard deviation for each trait, with green squares representing positive values and orange squares representing negative values, and hue depicting magnitude. Grey squares represent cases where no individual of a homozygous genotype was present in a treatment group.

**Supplementary Figure 6.**
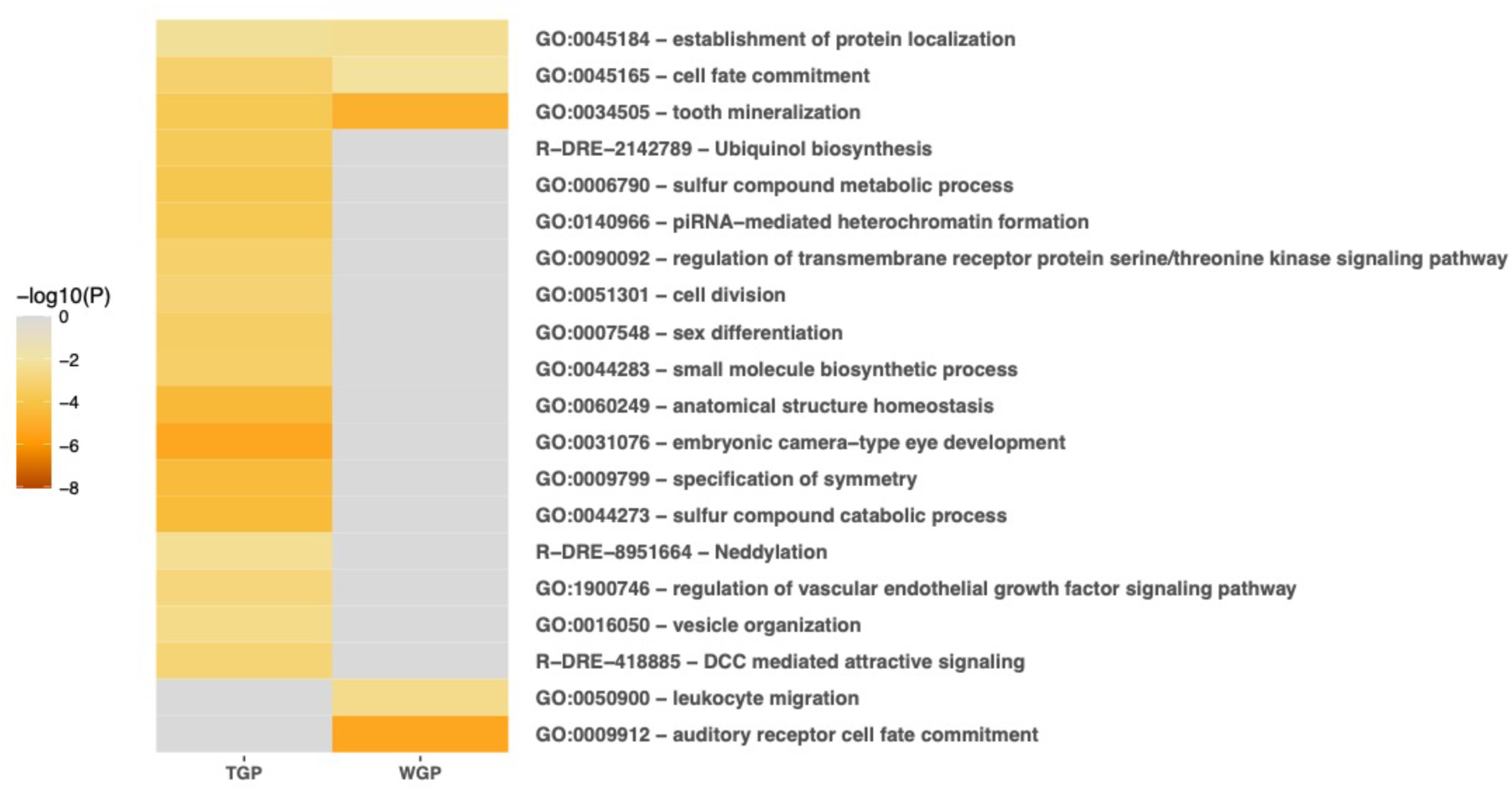
TGP and WGP patterns of methylation involve distinct biological processes. Top 20 terms enriched in candidate regions associated with **A)** Transgenerational (TGP) and **B)** Within-generation (WGP) patterns of methylation variation. Gene lists were submitted together and GO analysis was performed using metascape with *Danio rerio* gene IDs. Hue represents the -log10 p-value associated with each term, with darker colours representing lower p-values and hence greater statistical power.

## Supplementary Tables

**Supplementary Table 1.** Phenotypes used for QTL analysis and their respective trait groups.

| **Trait group** | **Trait** | **Calculation** |
|---|---|---|
| **Head Morphology** | Head shape | Derived from landmarks 1-12, 21-22, and semi-landmarks 23-32. First canonical root score for a *candisc* linear model identifying variation attributable to TGP or WGP respectively. |
|  | Eye Size | Centroid size of area between landmarks 3-5 |
|  | Size of pre-orbital region | Centroid size of area between landmarks 1-3 |
|  | Jaw Length | Linear Distance between landmarks 1 and 2 |
| **Body Morphology** | Body shape | As for ‘head shape’, but using landmarks 10-21 |
|  | Body depth | Linear distance between landmarks 12 and 21 |
|  | Size of caudal region | Centroid Size of area between landmarks 13 and 19 |
| **Methylation** | Methylation | LD1 of LDAs partitioning methylation variation based on F<sub>1</sub> and F<sub>2</sub> temperature respectively |

**Supplementary Table 2.** Genetic Map Parameters.

| **Chromosome** | **Number of Markers** | **Size (cM)** | **First marker** | **Last Marker** | **Genomic size (bp)** | **Coverage** |
|---|---|---|---|---|---|---|
| **1** | 396 | 91.995 | 2,048,648 | 25,893,570 | 29,619,991 | 80.53 |
| **2** | 504 | 107.724 | 21,992 | 23,681,314 | 23,686,546 | 99.58 |
| **3** | 365 | 225.129 | 66,228 | 17,625,865 | 17,759,012 | 98.87 |
| **4** | 598 | 122.439 | 1,726,372 | 32,975,342 | 34,181,212 | 91.42 |
| **5** | 885 | 110.520 | 103,965 | 15,544,997 | 15,550,311 | 99.29 |
| **6** | 571 | 147.762 | 142,322 | 17,959,476 | 18,825,451 | 94.64 |
| **7** | 894 | 116.583 | 99,321 | 30,503,803 | 30,776,923 | 98.78 |
| **8** | 901 | 118.909 | 21,796 | 20,469,523 | 20,553,084 | 99.48 |
| **9** | 826 | 144.433 | 319,604 | 20,795,074 | 20,843,631 | 98.23 |
| **10** | 539 | 167.552 | 53,939 | 14,481,084 | 17,985,176 | 80.22 |
| **11** | 446 | 170.223 | 685,814 | 16,053,532 | 17,651,971 | 87.05 |
| **12** | 565 | 130.665 | 26,518 | 19,018,170 | 20,694,444 | 91.77 |
| **13** | 500 | 127.358 | 44,166 | 20,466,720 | 20,748,428 | 98.43 |
| **14** | 382 | 166.803 | 19,381 | 15,833,815 | 16,147,532 | 97.93 |
| **15** | 457 | 174.073 | 58,112 | 16,909,641 | 17,318,724 | 97.30 |
| **16** | 427 | 182.813 | 121,162 | 16,554,909 | 19,507,025 | 84.24 |
| **17** | 357 | 78.151 | 2,417,704 | 19,845,618 | 20,195,758 | 86.29 |
| **18** | 287 | 115.539 | 313,489 | 14,497,577 | 15,939,407 | 88.98 |
| **19** | 445 | 162.781 | 1,576,694 | 20,487,073 | 20,580,295 | 91.88 |
| **20** | 656 | 156.629 | 60,564 | 20,340,378 | 20,445,003 | 99.19 |
| **21** | 341 | 163.910 | 563,109 | 15,307,182 | 17,421,465 | 84.63 |
| **Total** | 11,342 | 2981.991 |  |  |  | 92.79 |

**Supplementary Table 3.**
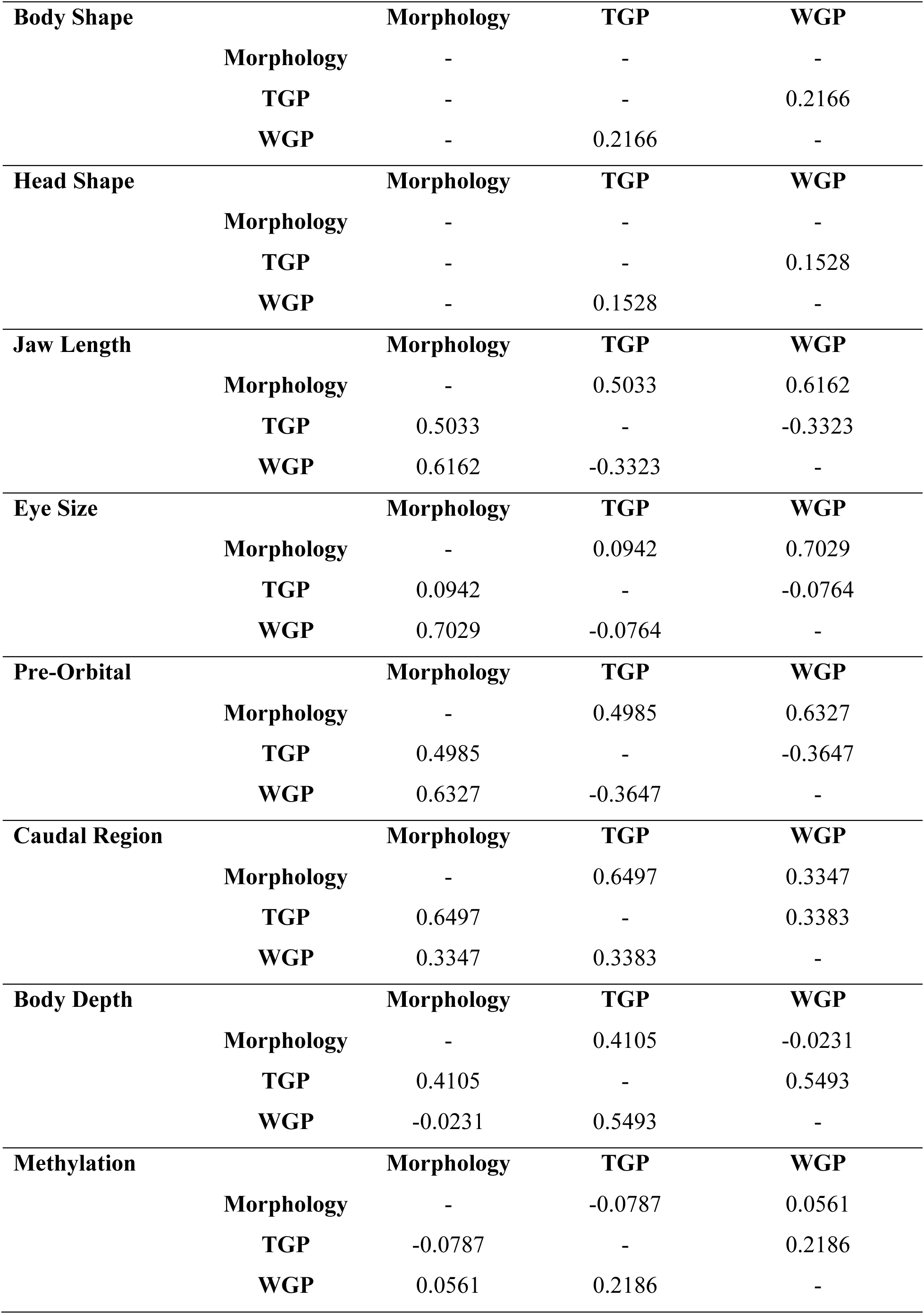
Partial pairwise correlation scores for each morphological trait.

| **Body Shape** |  | **Morphology** | **TGP** | **WGP** |
|---|---|---|---|---|
|  | **Morphology** | - | - | - |
|  | **TGP** | - | - | 0.2166 |
|  | **WGP** | - | 0.2166 | - |
| **Head Shape** |  | **Morphology** | **TGP** | **WGP** |
|  | **Morphology** | - | - | - |
|  | **TGP** | - | - | 0.1528 |
|  | **WGP** | - | 0.1528 | - |
| **Jaw Length** |  | **Morphology** | **TGP** | **WGP** |
|  | **Morphology** | - | 0.5033 | 0.6162 |
|  | **TGP** | 0.5033 | - | -0.3323 |
|  | **WGP** | 0.6162 | -0.3323 | - |
| **Eye Size** |  | **Morphology** | **TGP** | **WGP** |
|  | **Morphology** | - | 0.0942 | 0.7029 |
|  | **TGP** | 0.0942 | - | -0.0764 |
|  | **WGP** | 0.7029 | -0.0764 | - |
| **Pre-Orbital** |  | **Morphology** | **TGP** | **WGP** |
|  | **Morphology** | - | 0.4985 | 0.6327 |
|  | **TGP** | 0.4985 | - | -0.3647 |
|  | **WGP** | 0.6327 | -0.3647 | - |
| **Caudal Region** |  | **Morphology** | **TGP** | **WGP** |
|  | **Morphology** | - | 0.6497 | 0.3347 |
|  | **TGP** | 0.6497 | - | 0.3383 |
|  | **WGP** | 0.3347 | 0.3383 | - |
| **Body Depth** |  | **Morphology** | **TGP** | **WGP** |
|  | **Morphology** | - | 0.4105 | -0.0231 |
|  | **TGP** | 0.4105 | - | 0.5493 |
|  | **WGP** | -0.0231 | 0.5493 | - |
| **Methylation** |  | **Morphology** | **TGP** | **WGP** |
|  | **Morphology** | - | -0.0787 | 0.0561 |
|  | **TGP** | -0.0787 | - | 0.2186 |
|  | **WGP** | 0.0561 | 0.2186 | - |

**Supplementary Table 4.** Overlapping QTL between TGP and WGP models.

| **Chromosome** | **Trait** | **Model** | **Peak** | **CI start** | **CI end** |
|---|---|---|---|---|---|
| **4** | Body Depth | TGP | 27,254,817 | 15,201,200 | 30,439,939 |
|  | Body Depth | WGP | 28,870,398 | 14,029,739 | 30,439,939 |
| **5** | Head Shape | TGP | 8,920,391 | 8,829,631 | 11,876,249 |
|  | Methylation | TGP | 8,920,391 | 8,829,631 | 15,544,997 |
|  | Methylation | WGP | 15,435,758 | 7,830,756 | 15,544,997 |
|  | Jaw Length | WGP | 8,172,569 | 7,402,699 | 15,544,997 |
|  | Body Shape | WGP | 14,510,561 | 14,047,645 | 15,544,997 |
|  | Pre Orbital | WGP | 14,503,365 | 14,309,870 | 14,643,451 |
| **20** | Head Shape | TGP | 2,613,978 | 2,492,476 | 15,516,105 |
|  | Methylation | TGP | 15,414,990 | 12,536,388 | 16,410,200 |
|  | Pre_Orbital | TGP | 5,876,804 | 5,703,195 | 6,515,228 |
|  | Body_Shape | TGP | 2,613,978 | 2,492,476 | 19,525,672 |
|  | Caudal_Region | TGP | 13,393,375 | 7,490,101 | 20,340,378 |
|  | Caudal Region | WGP | 11,262,442 | 7,490,101 | 13,494,393 |

**Supplementary Table 5.** GO terms enriched in both TGP and WGP gene lists.

| **GO term** | **TGP logP** | **WGP logP** | **Best supported term in group** |
|---|---|---|---|
| GO:0048562: embryonic organ morphogenesis | -3.5094186 | -2.0364865 | GO:0048598: embryonic morphogenesis |
| GO:0034505: tooth mineralisation | -3.3770534 | -4.2958781 | GO:0034505: tooth mineralisation |
| GO:0001947: heart looping | -2.5600702 | -3.2324153 | GO:0048598: embryonic morphogenesis |
| GO:0003143; embryonic heart tube morphogenesis | -2.5826441 | -2.6266132 | GO:0048598: embryonic morphogenesis |
| GO:0051642: centrosome localization | -2.0175132 | -2.8690213 | GO:0051642: centrosome localization |
| GO:0006450: regulation of translational fidelity | -2.4247581 | -2.4739991 | GO:0006450: regulation of translational fidelity |
| GO:0018198: peptidyl-cysteine modification | -2.3372142 | -2.1012399 | GO:0018198: peptidyl-cysteine modification |

**Supplementary Table 6.**
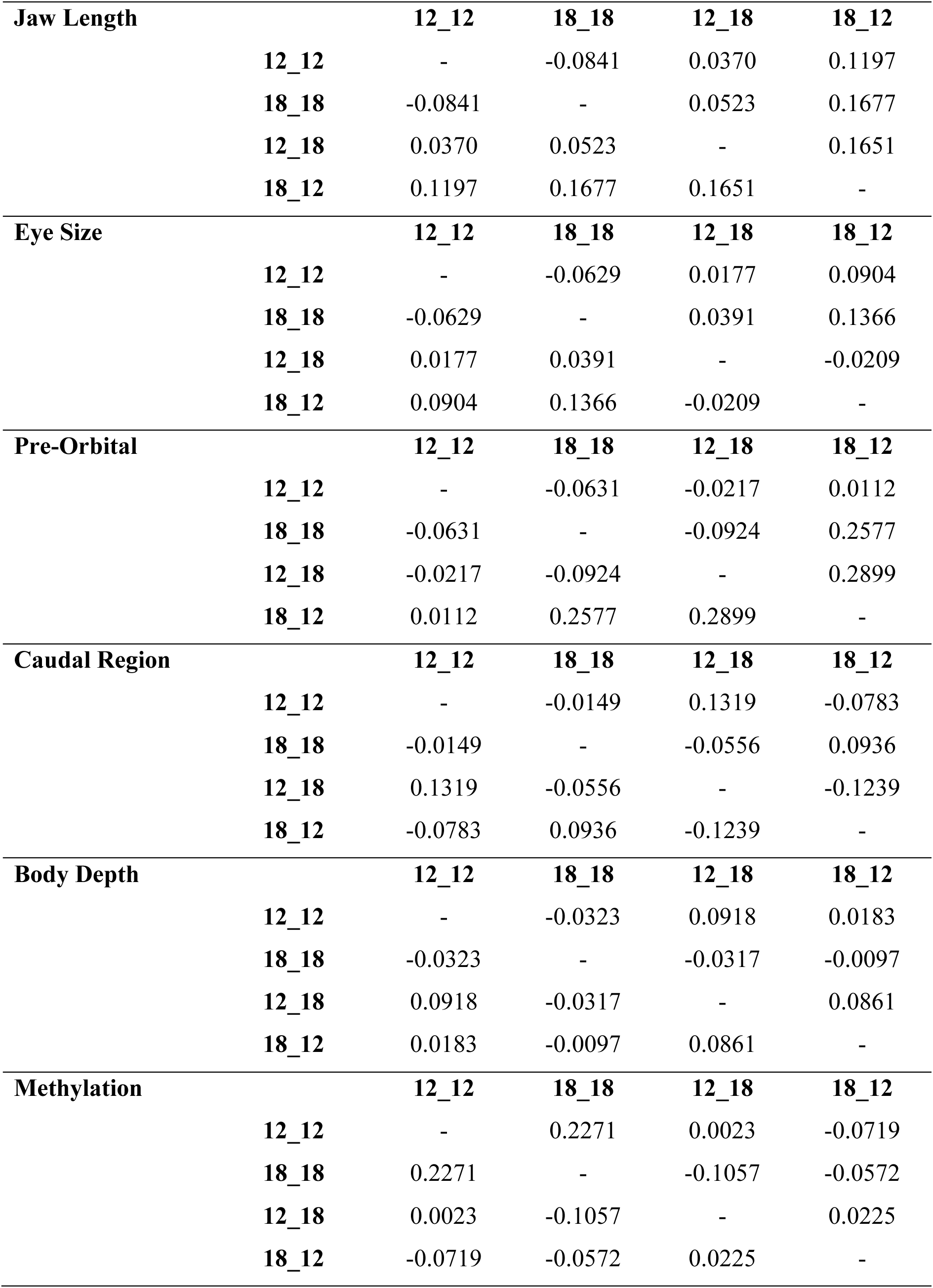
Partial pairwise correlation scores for each trait across multi-generational treatment groups.

